# The *R1441C-LRRK2* mutation induces myeloid immune cell exhaustion in an age- and sex-dependent manner

**DOI:** 10.1101/2023.10.12.562063

**Authors:** Rebecca L. Wallings, Karen McFarland, Hannah A. Staley, Noelle Neighbarger, Susen Schaake, Norbert Brüggemann, Simone Zittel, Tatiana Usnich, Christine Klein, Esther M Sammler, Malú Gámez Tansey

**Affiliations:** Department of Neuroscience, University of Florida, College of Medicine, McKnight Brain Institute, Gainesville, Florida, USA; Center for Translational Research in Neurodegenerative Disease, University of Florida, College of Medicine, McKnight Brain Institute, Gainesville, Florida, USA; Department of Neurology and Fixel Institute for Neurological Diseases, University of Florida Health, Gainesville, Florida, USA; Institute of Neurogenetics, University of Lübeck, Lübeck, Germany; Department of Neurology, University Medical Center Hamburg-Eppendorf, Hamburg, Germany; Aligning Science Across Parkinson’s (ASAP) Collaborative Research Network, Chevy Chase, MD, USA; MRC Protein Phosphorylation and Ubiquitylation Unit, School of Life Sciences, University of Dundee, Dundee DD1 5EH, U.K.; Department of Neurology, School of Medicine, Ninewells Hospital, Ninewells Drive, Dundee DD1 9SY, U.K

## Abstract

Considering age is the greatest risk factor for many neurodegenerative diseases, aging, in particular aging of the immune system, is the most underappreciated and understudied contributing factor in the neurodegeneration field. Genetic variation around the *LRRK2* gene affects risk of both familial and sporadic Parkinson’s disease (PD). The leucine-rich repeat kinase 2 (LRRK2) protein has been implicated in peripheral immune signaling, however, the effects of an aging immune system on LRRK2 function have been neglected to be considered. We demonstrate here that the *R1441C* mutation induces a hyper-responsive phenotype in macrophages from young female mice, characterized by increased effector functions, including stimulation-dependent antigen presentation, cytokine release, phagocytosis, and lysosomal function. This is followed by age-acquired immune cell exhaustion in a Lrrk2-kinase-dependent manner. Immune-exhausted macrophages exhibit suppressed antigen presentation and hypophagocytosis, which is also demonstrated in myeloid cells from *R1441C* and *Y1699C*-PD patients. Our novel findings that *LRRK2* mutations confer immunological advantage at a young age but may predispose the carrier to age-acquired immune cell exhaustion have significant implications for LRRK2 biology and therapeutic development. Indeed, LRRK2 has become an appealing target in PD, but our findings suggest that more research is required to understand the cell-type specific consequences and optimal timing of LRRK2-targeting therapeutics.

**One Sentence Summary:** The *R1441C*-*LRRK2* mutation causes an age-acquired immune cell exhaustion in macrophages in a sex-dependent manner

## INTRODUCTION

Parkinson’s disease (PD) is a common, progressive neurodegenerative disease, affecting around 1-2% of the population over the age of 65 (*1*). PD prevalence will likely increase two-fold by the year 2030 (*2*). In addition, the projected total economic burden will surpass $79 billion by 2037 (*3*), underscoring the need to reduce PD incidence and delay disease progression. The fact that there are currently no disease-modifying drugs for people with PD indicates that knowledge gaps still need to be closed to identify ways to cure or prevent this disease. Although the etiology of PD is largely unknown, it is thought to involve a complex interaction between various genetic and environmental factors (*4*). Although typically thought of as a disease limited to the central nervous system (CNS), evidence suggests a crucial and fundamental role of inflammation in both the CNS and the periphery in PD pathogenesis (*5*). Numerous studies of peripheral blood and cerebrospinal fluid from patients with PD suggest alterations in markers of inflammation and immune cell populations that could initiate or exacerbate neuroinflammation and perpetuate the neurodegenerative process (*6–9*). Furthermore, a number of disease genes and risk factors have been identified as modulators of immune function in PD (*10–12*).

Pathogenic variants in the gene encoding leucine-rich repeat kinase 2 (LRRK2) account for the majority of familial PD patients (*13*). LRRK2 is a large, complex protein containing six functional domains, two of which have known enzymatic activities; a ROC domain (GTPase activity) and a MAPKK domain (kinase activity). There are numerous single-nucleotide substitutions in each of these domains that have been shown to be pathogenic (*14–21*). The second most prevalent mutation, *R1441C*, located in the GTPase ROC domain, is present in about 46% of familial PD that has an origin in the Basque region in Spain (*22*) and leads to decreased LRRK2 GTPase hydrolysis and increased LRRK2 protein kinase activity (*23, 24*).

LRRK2 is expressed in innate and adaptive immune cells and its expression is tightly regulated by immune stimulation (*9, 25–28*). Furthermore, LRRK2 is a member of the receptor interacting protein (RIP) kinase family, which is a group of proteins that detect and respond to cellular stress by regulating cell death and activation of the immune system (*29*). LRRK2 function has also been implicated in pathogen control based on observations that LRRK2 variants confer differential susceptibility to a number of bacterial infections, including *Mycobacterium tuberculosis* (*30, 31*), *Salmonella Typhimurium* (*32*), and *Listeria monocytogenes* (*33*), as well as an increased risk of developing leprosy (*34*) and inflammatory bowel disease such as Crohn’s (*35*) and ulcerative colitis (*36*). Collectively such data strongly suggest a generalized role of LRRK2 in immune system regulation.

Although research has focused primarily on neuroinflammation in PD over the last two decades, it is becoming increasingly clear that peripheral inflammatory responses may play important roles in PD pathogenesis (*37, 38*). For example, immune signaling plays a role in the regulation of neurodegeneration in a *LRRK*2-PD pre-clinical model, with the *R1441C* mutation altering the type II interferon response in peripheral immune cells (*28*). Furthermore, neutralization of IL-6 over-production in the periphery of these mice prevents substantia nigra *pars compacta* (SNpc) dopaminergic (DA) neuron loss in the presence of peripheral lipopolysaccharide (LPS) exposure (*39*). Importantly, immune dysfunction in the periphery takes primacy over the CNS, as the presence of *Lrrk2* mutations in lymphocytes alone is sufficient to induce DA neuron loss in the midbrain of LPS-treated mice (*39*). Such data suggest a pertinent role of the peripheral immune system in *LRRK2*-PD, with immune dysfunction driving the development of the disease.

As well as a role in immune response, a vast amount of research has focused on the role of LRRK2 at the lysosome. In immune cells, specifically, LRRK2 has been shown to mediate a number of lysosome-dependent immune functions such as phagocytosis (*40*) and antigen presentation (*9, 27, 41*). Intriguingly, LRRK2 has been shown to alter lysosomal function and autophagy in kidneys in an age-dependent, biphasic manner, with enhanced autophagy levels in young *Lrrk2*-null mice, that are reduced and drop below control levels once the mice reach 20 months of age (*42*). Despite this intriguing observation, few studies have investigated the effects of *LRRK2* mutations on lysosomal processes in the context of aging. Considering age is the greatest risk factor for many neurodegenerative diseases, aging, in particular aging of the immune system (*5*), is the most underappreciated and understudied contributing factor to neurodegeneration in the field.

To close this important gap, we assessed the effects of *LRRK2* mutations on immune responses and lysosomal function in peripheral immune cells in young versus old mice. We used *R1441C-Lrrk2* mutant knock-in (KI) mice based on previous findings demonstrating an effect of the *R1441C/G* mutation on peripheral immune responses (*28, 39*) whilst evading limitations of an overexpression model and investigating the effects of the *R1441C* mutation in physiologically relevant levels. Peritoneal macrophages (pMacs) were isolated from these mice to examine the effects of the *R1441C* mutation on macrophage cell function at different ages. pMacs are not a homogenous population of cells, but rather a mix of small and large pMacs (SPMs and LPMs, respectively). LPMs are resident to the peritoneal cavity and are traditionally thought of as anti-inflammatory, phagocytic and responsible for the presentation of exogenous antigens (*43*). SPMs, on the other hand, are generated from bone-marrow-derived myeloid precursors which migrate to the peritoneal cavity in response to infection, inflammatory stimuli, or thioglycolate, and present a pro-inflammatory functional profile (*43*). We observed a bi-phasic, age-dependent effect of the *R1441C-Lrrk2* KI mice, with increased antigen presentation, production of anti-inflammatory cytokines, lysosomal activity, and pathogen uptake in peripheral macrophages from young (2-3 months) *R1441C-Lrrk2* KI mice examined *ex vivo*. Conversely, peripheral macrophages from aged (18-21 months) *R1441C-Lrrk2* KI mice exhibited a diminished capacity to present antigens in the presence of inflammatory insults, decreased lysosomal function and pathogen uptake, with concomitant increases in DNA fragmentation in the presence of pathogens. Importantly, we demonstrate that this is a sex-dependent phenotype, with male *R1441C-Lrrk2* KI failing to present this age-dependent phenotype. Furthermore, we demonstrate that the age-dependent hypophagocytosis and decreased antigen presentation in the *R1441C-LRRK2* KI females is due to an acquired immune cell exhaustion driven by increased Lrrk2 protein kinase activity and immune responses whilst young. These phenotypes were also observed in human peripheral myeloid cells, with monocyte-derived macrophages from *R1441C* and *Y1699C* patients exhibiting decreased pathogen uptake and increased programmed cell death protein ligand 1 (PDL1) expression, consistent with immune cell exhaustion.

## RESULTS

### *LRRK2 R1441C* mutant immune cells display age-dependent biphasic alteration in antigen presentation

LRRK2 protein expression has been shown to increase in immune cells from both murine pre-clinical models and patient cells in response to inflammatory stimuli (*9, 27, 28, 44–46*). To determine whether this is age-dependent, Lrrk2 protein expression were assessed in pMacs from young and aged mice treated with 100U of IFNγ for 18h. In pMacs from young females, IFNγ treatment significantly increased total Lrrk2 protein in both B6 and *R1441C* pMacs (Fig. 1A, B; p < 0.01). Relative to pMacs from young female mice, pMacs from aged females expressed increased total Lrrk2 protein under vehicle stimulation, with aged *R1441C* pMacs expressing significantly less Lrrk2 protein relative to B6 aged pMacs (p <0.05). Furthermore, pMacs from aged females did not exhibit a significant increase in Lrrk2 protein expression with IFNγ treatment, with aged *R1441C* pMacs still expressing significantly less Lrrk2 protein relative to B6 aged pMacs in IFNγ-treatment (p<0.05). In pMacs from young male mice, the same observations were made as in the females (Fig. S1A, B). In aged male mice, although a lack of IFNγ-mediated Lrrk2 increase was observed as seen in females, the effect of the *R1441C* mutation was not observed. These observations indicate that Lrrk2 protein increase in macrophages as a function of age in both males and females, however, the *R1441C* mutation limits this increase in a sex-dependent manner.

**Figure 1.**
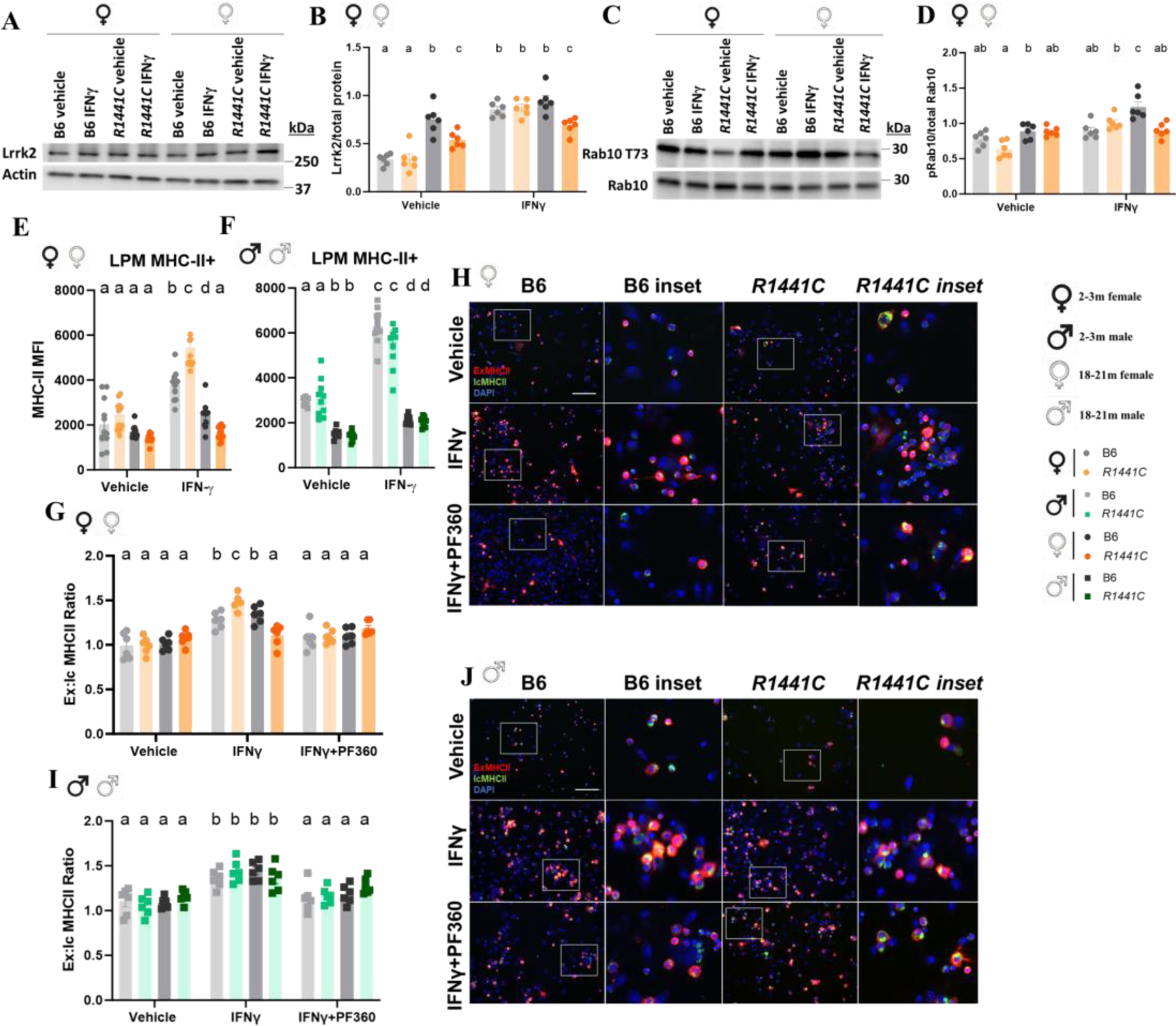
*R1441C* mutation leads to age dependent biphasic alteration in antigen presentation. pMacs from 2-3-month and 18-21-month-old female or male, B6 or *R1441C* mice were stimulated with 100U IFNγ for 18-hours. (A, B) Total Lrrk2 protein expression was assessed and normalized to β-Actin and quantified. Representative western blots shown. (C, D) T73 Rab10 protein expression was assessed and normalized to total Rab10 and quantified. Representative western blots shown. (E, F) Surface MHC-II median fluorescence intensity (MFI) was quantified on LPMs and via flow cytometry. (G-J) pMacs were stained for intracellular and extracellular MHC-II MFI and Ex:IcMHC-II ratio quantified. Scale bars, 100μM. Bars represent mean +/- SEM (N = 6-12 mice). Three-way ANOVA, Bonferroni *post hoc*, groups sharing the same letters are not significantly different (p > 0.05) whilst groups displaying different letters are significantly different (p < 0.05).

To decipher whether these alterations in Lrrk2 protein expression was accompanied by increases in Lrrk2 kinase activity, phosphorylation of Rab10 at T73 (pT73), a *bona fide* kinase-substrate of Lrrk2 (*23*), was quantified. In pMacs from young female mice, a significant (p<0.01) increase in pT73 Rab10 expression was observed in response to IFNγ in *R1441C-Lrrk2* cells, whilst this was not observed in pMacs from young B6 females (Fig. 1C, D). Interestingly, this phenotype was reversed in pMacs from aged female mice, with a significant increase (p<0.01) in pT73 Rab10 expression observed in response to IFNγ in B6 but not *R1441C*-Lrrk2 cells. Of note, the pT73 Rab10 expression in pMacs from aged B6 females in response to IFNγ was significantly (p<0.05) elevated relative to all other groups, suggesting that, in B6 females, Lrrk2 kinase activity increases with age. In pMacs from young male mice, however, no significant effect of the *R1441C* mutation was observed in vehicle or IFNγ conditions (Fig. SB, C), although a significant (p<0.01) increase in pT73 Rab10 expression was observed in response to IFNγ. Interestingly, a significant increase in pT73 Rab10 expression was observed in pMacs from aged male mice relative to those from young males, suggesting that Lrrk2 kinase activity increases with age in pMacs from male mice. Furthermore, upon IFNγ treatment, a significant increase in pT73 Rab10 expression was observed in pMacs from aged *R1441C-Lrrk2* males relative to all other groups, suggesting an additive effect of the *R1441C* mutation on Lrrk2 kinase activity in an ageing male immune system.

As pMacs are not a homogenous population of cells, as previously discussed, we used multi-color flow cytometry-based methods to immunophenotype LPMs and SPMs, as the two populations can be distinguished based on CD11b expression (Fig. S1D) (*43*). No differences were seen in LPM or SPM count between genotypes in males or females at either young or aged (Fig. S1E-H).

To begin to investigate the effects of the *R1441C* mutation on pMac function at different ages, surface MHC-II expression was assessed on both LPMs and SPMs via multi-color flow cytometry as a measure of antigen presentation in both vehicle and IFNγ treated pMacs. In pMacs from young females, the *R1441C* mutation increases surface MHC-II expression on LPMs in IFNγ treatments (Fig 1D). Interestingly, although no effect of age was observed in vehicle conditions, surface MHC-II expression was significantly decreased in aged pMacs from both B6 and *R1441C* when treated with IFNγ relative to LPM from young mice (p<0.001). Furthermore, a significant decrease in surface MHC-II expression was observed in LPM from aged *R1441C* females relative to aged B6 LPM (p<0.01), indicative of decreased ability to engage in antigen presentation in these cells. This effect of the *R1441C* mutation was not observed in LPMs from male mice (Fig. 1E). However, as seen in LPM from female mice, an effect of age was also observed in LPM from male mice, with surface MHC-II expression significantly decreased in LPM from aged mice, in both vehicle and IFNγ conditions (p<0.001). Regarding surface MHC-II expression on SPMs, the same effect of age was observed in both male and female pMacs, with SPM from aged mice expressing significantly less surface MHC-II relative to SPM from young mice (p<0.001) (Fig. S1I, J).

With alterations in surface MHC-II expression, there is likely increased and decreased transport of MHC-II-complexes to the cell surface in pMacs from young and aged *R1441C* females, respectively. To test this, using pMacs treated with vehicle or IFNγ, we co-stained for extracellular vs intracellular MHC-II (ExMHCII vs IcMHCII) and monitored expression via fluorescence microscopy. pMacs were co-treated with 100nM PF360 to determine the effects of Lrrk2 kinase activity on surface MHC-II expression. Treatment with the Lrrk2 kinase inhibitor, PF360 (PF-06685360)PF360 reduced expression of Lrrk2 phosphorylated at S935 (pS935) Lrrk2 in all groups (Fig. S2A, B). Although no significant effects of genotype or age were observed in vehicle treated pMacs from female mice, upon IFNγ stimulation, a significant increase in Ex:IcMHCII ratio was observed in pMacs from young *R1441C* mice relative to other groups (p<0.05) (Fig. 1F, Fig. S2C). Furthermore, a significant decrease in Ex:IcMHCII ratio was observed in pMacs from aged *R1441C* females relative to aged B6 pMacs and young pMacs (p<0.01) (Fig. 1G). The increased Ex:IcMHCII ratio in pMacs from young *R1441C* females was ameliorated upon Lrrk2 kinase inhibition (Fig. S2C, D), with this treatment also decreasing Ex:IcMHCII ratio in pMacs from young and aged B6 females. However, no effect of Lrrk2 kinase inhibition was seen in pMacs from aged *R1441C* females. Such data suggests that Lrrk2-kinase inhibition reduces the transport of IcMHCII to the cell surface to engage in antigen presentation, but this may occur in an age- and genotype-dependent manner. No effect of the *R1441C* mutation was observed in males (Fig. 1H, I, Fig. S2D). It seems therefore that the *R1441C* mutation exhibits a bi-phasic, age- and sex-dependent effect on antigen presentation in macrophages. The effect of Lrrk2 protein kinase inhibition on antigen presentation, however, does not appear to be bi-phasic, with Lrrk2 kinase inhibition decreasing antigen presentation in pMacs from both young and aged mice, at least in B6 mice.

### *LRRK2 R1441C* mutant immune cells display age-dependent biphasic alteration in cytokine release ex vivo and circulating cytokines in vivo

To investigate the extent to which alterations in antigen presentation were accompanied by changes in cytokine release, media from vehicle- and IFNγ treated cells were collected and cytokine release was quantified on a multiplexed immunoassay panel (Fig. 2A). In pMacs from young *R1441C* females, increases in IL6 relative to B6 controls was observed in vehicle treated conditions, which was ameliorated upon ageing (Fig. 2B). Interestingly, a bi-phasic, age-dependent pattern was observed regarding IL10 release, with increased IL10 observed in pMacs from young *R1441C* females relative to young B6 controls, whilst, conversely, decreased IL10 was observed in pMacs from aged *R1441C* females relative to aged B6 controls (Fig. 2B). As well, TNF was significantly elevated in IFNγ-treated pMacs from R1441C aged females relative to aged B6 controls and both young B6 and young *R1441C* females (p<0.05) (Fig. 2B). With a decrease in the release of the anti-inflammatory cytokine, IL10, and an increase in the release of the pro-inflammatory cytokine, TNF, in females, these findings indicate that pMacs from aged *R1441C* female mice promote a pro-inflammatory microenvironment.

**Figure 2.**
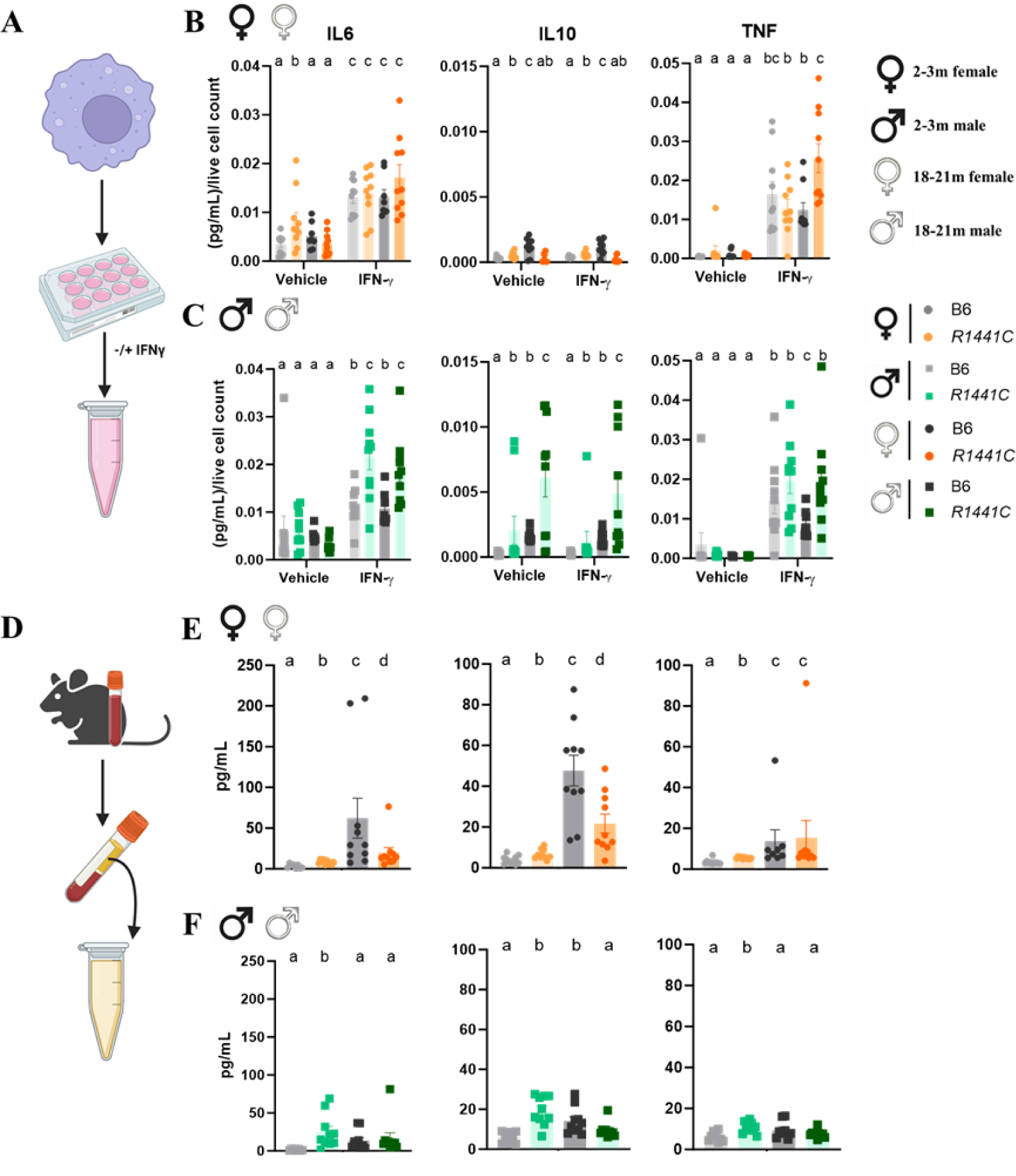
*R1441C* mutation leads to age dependent biphasic alteration in cytokine release ex vivo and circulating cytokines in vivo. (A) pMacs from 2-3-month and 18-21-month-old female or male, B6 or *R1441C* mice were stimulated with 100U IFNγ for 18-hours and media collected. (B, C) IL10, IL6 and TNF were quantified and normalized to live cell count. (D) Trunk blood was collected via trunk bleed and plasma was isolated. (E, F) IL10, IL6 and TNF were quantified. Bars represent mean +/- SEM (N = 8-10 mice). Three-way ANOVA, Bonferroni *post hoc*.

In pMacs from *R1441C* males, IL6 release was significantly increased in IFNγ conditions relative to B6 controls in both young and aged mice (p<0.05) (Fig. 2C). Similarly, IL10 release was significantly increased in pMacs from both aged and young *R1441C* males relative to B6 controls (p<0.05) (Fig.2C). Regarding TNF release from pMacs from male mice, no genotype effects were observed in pMacs from young mice, whilst an increase in TNF was observed in aged R1441C males relative to aged B6 controls, although this did not significantly differ from pMacs from neither young B6 or *R1441C* males (Fig.2C). It seems therefore that, in pMacs from aged male mice, the *R1441C*-associated increase in IL6 and IL10 seen in young mice is retained with age but *R1441C* macrophages secrete more TNF than B6 controls after stimulation.

To determine if these alterations in cytokine release associated with the *R1441C* mutation *ex vivo* was also detectable *in vivo*, plasma from whole blood was collected and cytokine release was quantified (Fig. 2D). A significant increase in IL10, IL6, and TNF was observed in plasma from young *R1441C* female mice relative to young B6 controls (p<0.05) (Fig. 2E). Interestingly, an increase in these cytokines were observed with age, although IL6 and IL10 release was significantly reduced in aged *R1441C* females relative to B6 controls (p<0.01), confirming the biphasic age-dependent effects observed *ex vivo*. In young male mice, a significant increase in both IL6, IL10 and TNF were also observed in *R1441C* males relative to B6 controls (p<0.01) (Fig. 2F). However, unlike in females, an age-dependent increase in cytokine release across genotypes was not observed, with the only genotype effect being observed in aged male mice seen with IL10, with a decrease in IL10 observed in aged *R1441C* males relative to aged B6 controls.

### The R1441C Lrrk2 mutation is associated with alterations of monocyte populations and microglia activation in aged female mice

The observation of a biphasic immune response, with an initial hyperinflammation state proceeded by a subsequent downregulation of immune responses, prompted us to hypothesize that what was being observed in aged *R1441C* female immune cells was immune cell exhaustion. Immune cell exhaustion arises from chronic stimulation of immune cells, leading to a downregulation, or even absence, of an expected effector response (e.g. antigen presentation, cytokine release or phagocytosis) (*47*). It is known that monocytes pushed to exhaustion shift to express higher Ly6C levels and a reduction in Ly6C^-^ monocytes under resting conditions (*48*). We therefore assessed monocyte sub-populations in the blood of young and aged mice to determine if a shift to an exhausted state could be observed. Indeed, we observed a detectable increase in Ly6C^+^:Ly6C^-^ in the peripheral blood mononuclear cells isolated from blood of aged *R1441C* female mice relative to aged B6 controls and young B6 and *R1441C* female mice (Fig. S3A) with no significant change observed in male mice (Fig. S3B). The fact that no effect of genotype was observed in young females and only aged females, suggests the development of an age-acquired immune cell exhaustion phenotype in the *R1441C* females. This is further supported by the observation that in aged *R1441C* female mice, but not males, an increase in PDL1 expression, an immune checkpoint molecule, was upregulated on LyC+ monocytes from blood relative to B6 controls (Fig. S3C, D).

It has been suggested that circulating monocytes, disrupted in PD, may infiltrate the brain parenchyma and affect local glial and adaptive immune responses (*49*). To determine whether exhausted monocytes could be infiltrating the brain parenchyma of aged *R1441C* female mice, immune cells were isolated from whole brain and assessed via multi-color flow cytometry (Fig. S3E). Indeed, in aged *R1441C* females, an increase in overall Ly6C^+^ monocytes in the brain parenchyma was observed relative to B6 controls, whilst MHC-II/Ly6C^+^ were decreased (Fig. S3F, G), suggesting that whilst there is an increase in infiltrating Ly6C^+^ monocytes in the brains of aged *R1441C* females, they exhibit the same decrease in antigen presenting capabilities to those observed in pMacs *ex vivo*. Importantly, alongside this increase in infiltrating monocytes into the brain parenchyma, a concomitant increase in MHC-II expression on microglia was observed in aged *R1441C* females relative to B6 controls (Fig. S3H) with no changes in overall microglia number (Fig. S3I). No changes were observed in the brains of male mice (Fig. S3J-M).

### *LRRK2 R1441C* mutant immune cells display age-dependent biphasic alterations in lysosomal activity

As previously discussed, loss of *Lrrk2* leads to biphasic alterations in lysosomal function in kidneys (*42*). Interestingly, lysosomal dysfunction has been implicated in immune cell exhaustion, with decreased autophagy levels and lysosomal function being associated with increased T cell exhaustion (*50*). We therefore hypothesized that the biphasic immune response observed in *R1441C* females would be associated with concomitant biphasic lysosomal function.

The Eα: YAe model is used to monitor the antigen presentation capabilities of cells by incubating an endogenous peptide (Eα 52-68) which is subsequently phagocytosed, transported to the lysosome, loaded onto an MHC-II complex at the lysosome and transported back to the plasma membrane for antigen presentation (Fig. 3A). This Eα peptide-loaded MHC-II can subsequently be detected via flow cytometry using the YAe antibody (*51*); DOI: 10.1016/j.jim.2015.04.023). We observed that YAe median fluorescence intensity (MFI) was significantly increased in IFNγ-treated LPMs from young *R1441C* females relative to age-matched B6 controls and pMacs from aged females (p<0.05) (Fig. 3B). Furthermore, IFNγ-treated LPMs from aged *R1441C* females exhibited decreased YAe MFI relative to other groups. Co-treatment with PF360 ameliorated the increased YAe MFI observed in LPMs from young *R1441C* females. Antigen presentation and pathogen sensing require protease action and sufficient lysosomal function in order to occur (*52*), which is why lysosomotropic agents have been shown to decrease peptide-loaded-MHC-II surface expression in antigen presenting cells (*51*). Indeed, when co-treated with the vacuolar H+ ATPase (V-ATPase) inhibitor, Bafilomycin A1, YAe MFI significantly decreased in both *R1441C* and B6 LPMs from both young and aged mice (p<0.01). Although no effect of genotype was observed in male mice at either ages, the same treatment effects were observed as seen in females, with PF360 and Bafilomycin A1 co-treatment significant decreasing YAe MFI relative to IFNγ conditions (p<0.01) (Fig. 3C). Due to this observation and the crucial role of the lysosome in antigen presentation, we next sought to probe the effects of the *R1441C* mutation on lysosomal function in these pMacs.

**Figure 3.**
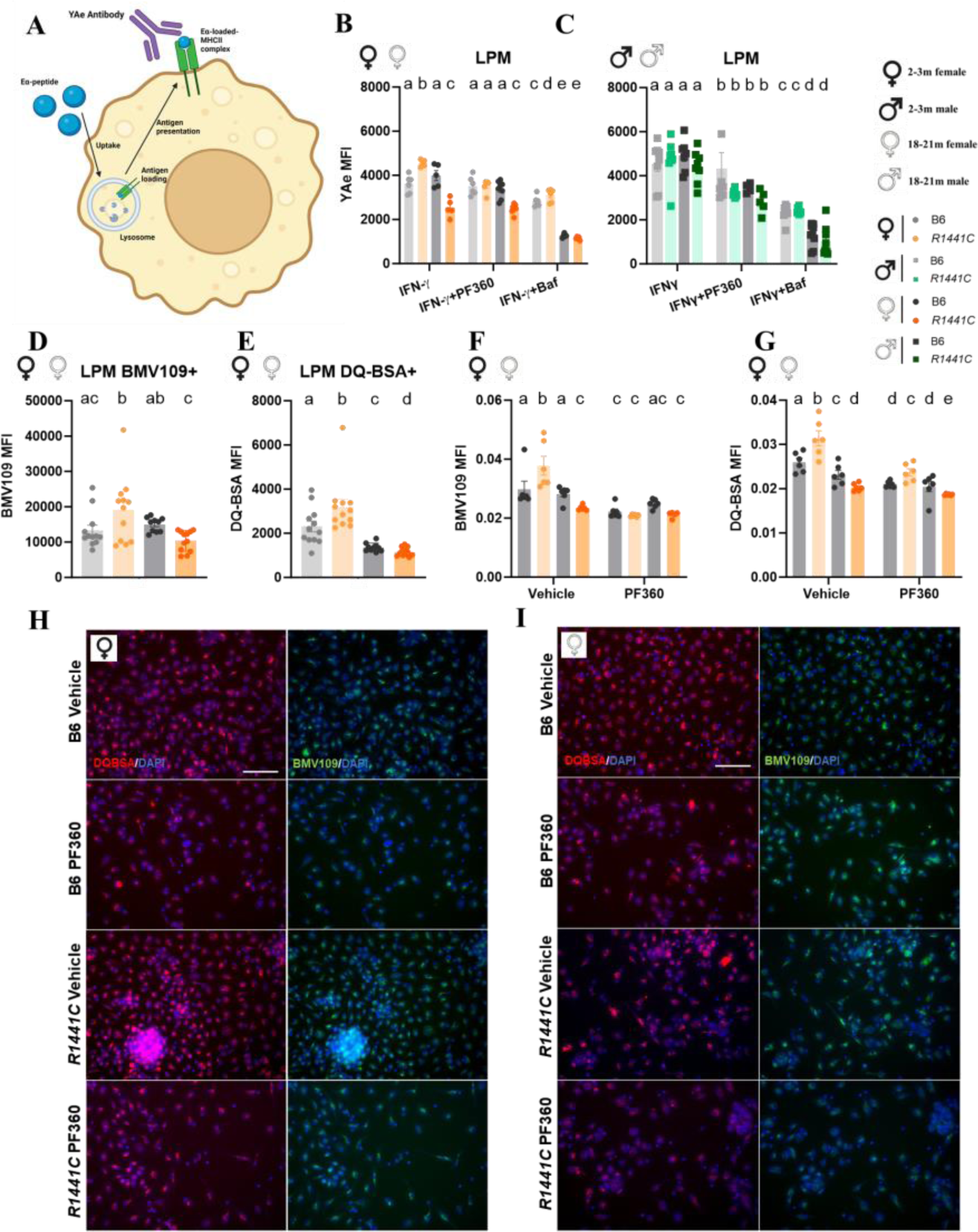
*R1441C* mutation leads to age dependent biphasic alteration in lysosomal activity. pMacs from 2-3-month and 18-21-month-old female or male, B6 or *R1441C* mice were stimulated with 100U IFNγ for 18-hours. (A) Schematic of YAe assay. (B, C) Cells were co-treated with 100nM PF360 or 50nM Bafilomycin A1 (baf) and YAe MFI was quantified on LPMs via flow-cytometry. (D, E) DQ-BSA and BMV109 MFI was quantified in LPMs via flow cytometry. (F-I) DQ-BSA and BMV109 MFI was quantified in pMacs via microscopy. Scale bars, 100μM. Bars represent mean +/- SEM (N = 6-12 mice). Two -way ANOVA, Bonferroni *post hoc*, or Student’s T-test. Groups sharing the same letters are not significantly different (p > 0.05) whilst groups displaying different letters are significantly different (p < 0.05).

pMacs were plated and incubated with the lysosomal degradation probe, DQ Red BSA, and the pan-cathepsin probe, BMV109, and collected for flow cytometry to quantify lysosomal activity. It was observed that LPMs from young *R1441C* females exhibited increased DQ-BSA and BMV109 MFI relative to age-matched controls and pMacs from aged B6 and *R1441C* females (Fig. 3D, E), indicative of increased lysosomal protein degradation and pan-cathepsin activity, respectively. Conversely, LPMs from aged *R1441C* females exhibited decreased DQ-BSA and BMV109 MFI relative to age-matched controls and pMacs from young B6 and *R1441C* females. Interestingly, LPM from both aged B6 and *R1441C* females exhibited decreased DQ-BSA MFI relative to young mice, reflecting an overall decrease in lysosomal degradative capacity with age. No effects of genotype were observed in LPMs from male mice (Fig. S4A, B). These observations made via flow cytometry were confirmed with fluorescence microscopy (Fig. 3F-I; Fig. S4C-F). As was observed with YAe measurements, Lrrk2 kinase inhibition via treatment with PF360 significantly reduced BMV109 MFI observed via microscopy (p<0.05) in all groups except aged B6 and *R1441C* females, which exhibited decreased measurement of pan cathepsin activity which were unaffected by Lrrk2 kinase inhibition. However, Lrrk2 kinase inhibition was able to decrease DQ-BSA MFI in all groups.

### *R1441C* Lrrk2 mutation is association with age-acquired hypo-phagocytosis

Lysosomal function in macrophages is crucial for a number of other immune-related functions in myeloid cells, including phagocytosis. Therefore, we posited that the observed biphasic effects of the *R1441C* mutation on lysosomal function would have downstream consequences on phagocytosis in pMacs. To explore this hypothesis, pMacs were plated and incubated with pHrodo™ Green *E. coli* BioParticles™, after which uptake and fluorescence intensity were monitored over 5 hours. An increase in GFP MFI was observed in pMac cultures from young *R1441C* females relative to age-matched B6 controls and pMac cultures from aged B6 and *R1441C* females (Fig. 4A, B). As well, a significant reduction in GFP MFI was observed in pMacs from aged *R1441C* females relative to age-matched B6 controls and young B6 and *R1441C* females (p<0.01). Lrrk2 kinase inhibition with PF360 significantly reduced GFP MFI in young *R1441C fe*males (p<0.01), with no effects of Lrrk2 kinase inhibition on GFP MFI in other groups. In male mice, pMacs from young *R1441C* males did exhibit increased GFP MFI relative to age-matched B6 controls and aged B6 and *R1441C* mice which could be significantly reduced with PF360 co-treatment (p<0.05) (Fig. S5A-B). Interestingly, an overall effect of age was observed in pMac cultures from male mice, with aged B6 and *R1441C* pMacs exhibiting decreased GFP MFI relative to pMacs cultures from young male mice. It was still observed, however, that, despite this decrease in phagocytosis with age, pMacs from aged *R1441C* males exhibited increased GFP MFI relative to age-matched B6 controls. Collectively, it seems that pMacs from *R1441C* females exhibit increased phagocytic uptake at a young age and develop an age-dependent hypophagocytic phenotype. pMacs from *R1441C* males, however, exhibit this same increased phagocytic uptake at a young age but retain this increase in old ages.

**Figure 4.**
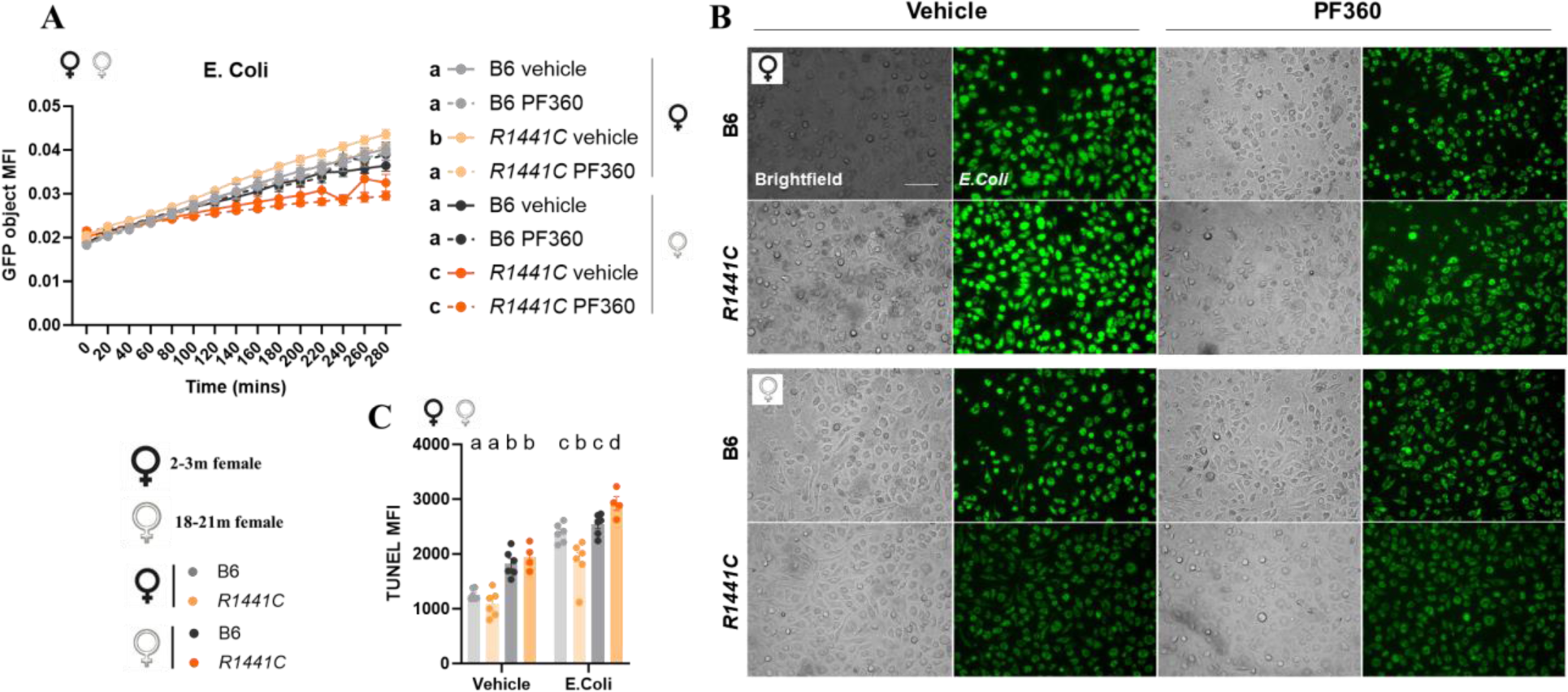
*R1441C* Lrrk2 mutation alters pathogen uptake and pathogen-mediated DNA damage in age- and sex-dependent manner. pMacs from 2-3-month and 18-21-month-old female, B6 or *R1441C* mice were incubated with 100μg of pHrodo *E. coli*. +/- 100nM PF360. (A, B) pMacs were imaged every 20 minutes for 5 hours, and GFP fluorescence was measured. Scale bars, 100μM. Four-way ANOVA, Bonferroni *post hoc*. Main effects of groups are reported in the graph key. Groups sharing the same letters are not significantly different (p > 0.05) whilst groups displaying different letters are significantly different (p < 0.05). (C) TUNEL MFI was quantified in LPMs via flow cytometry. Bars represent mean +/- SEM (N = 4-12 mice). One-way ANOVA, Bonferroni *post hoc*. Groups sharing the same letters are not significantly different (p > 0.05) whilst groups displaying different letters are significantly different (p < 0.05).

### Hypophagocytic activity in *R1441C* macrophages is associated with aberrant pathogen-mediated DNA damage

Our data thus far suggests a potential age-acquired immune cell exhaustion in macrophages from *R1441C* female mice. In other disease states associated with cellular exhaustion, apoptotic depletion of immune cells is also observed (*53*). To investigate the degree of pathogen-induced apoptosis in pMacs from *R1441C* mice, pMacs were plated in the presence of *E. coli* for 12 hours, harvested and stained using terminal deoxynucleotidyl transferase dUTP nick end labeling (TUNEL), which detects DNA breaks formed during the final phase of apoptosis, and TUNEL fluorescence was assessed via flow cytometry. LPMs from young *R1441C* females exhibited decreased TUNEL MFI in response to *E. Coli* relative to age-matched B6 controls and LPMs from aged mice, whilst an increase was observed in LPMs from aged *R1441C* females relative to all other groups (Fig. 4C). Reduced apoptosis in young *R1441C* mice could lead to prolonged presence of activated immune cells which should otherwise be eliminated, resulting in an increased and prolonged inflammatory response and, if sustained for prolonged times, could lead to immune cell exhaustion. Indeed, in response to *E. Coli*, no effect of genotype was observed in aged males (Fig. S5C). These novel findings support the conclusion that *R1441C* males may evade the age-acquired immune cell exhaustion that is seen in females by increasing clearance of activated immune cells.

### Immune cell exhaustion in *R1441C* Lrrk2 mutant immune cells is dependent on Lrrk2 kinase activity

Our data thus far suggest a potential age-acquired immune cell exhaustion in macrophages from *R1441C* female mice. We hypothesized that this age-acquired immune cell exhaustion was driven by the initial hyperinflammatory phase when these mice are young, characterized by increased cytokine release, increased antigen presentation, and reduced apoptosis of activated immune cells. We therefore hypothesized that pMacs from young *R1441C* female mice could be pushed to acquire this age-dependent immune cell exhaustion phenotype if subjected to chronic overstimulation *ex vivo.* To test this, pMacs were isolated and plated in the presence of a vehicle or 100ng/mL LPS for a 5-day chronic treatment (Fig. 5A). We first sought to understand how Lrrk2 expression is altered by LPS-induced exhaustion and how age and the *R1441C* affects this. In pMacs from young females, an increase in Lrrk2 expression was observed in response to LPS-induced exhaustion (Fig. 5B, C). Importantly, there was no significant effect of treatment in pMacs from aged female mice, with aged *R1441C* pMacs expressing significantly less Lrrk2 relative to B6 controls (p<0.01). Interestingly, these same observations were made in pMacs from aged male mice (Fig. S6A, B) suggesting that, with an additional chronic inflammatory stress, aged *R1441C* males can exhibit some of the phenotypes observed in females.

**Figure 5.**
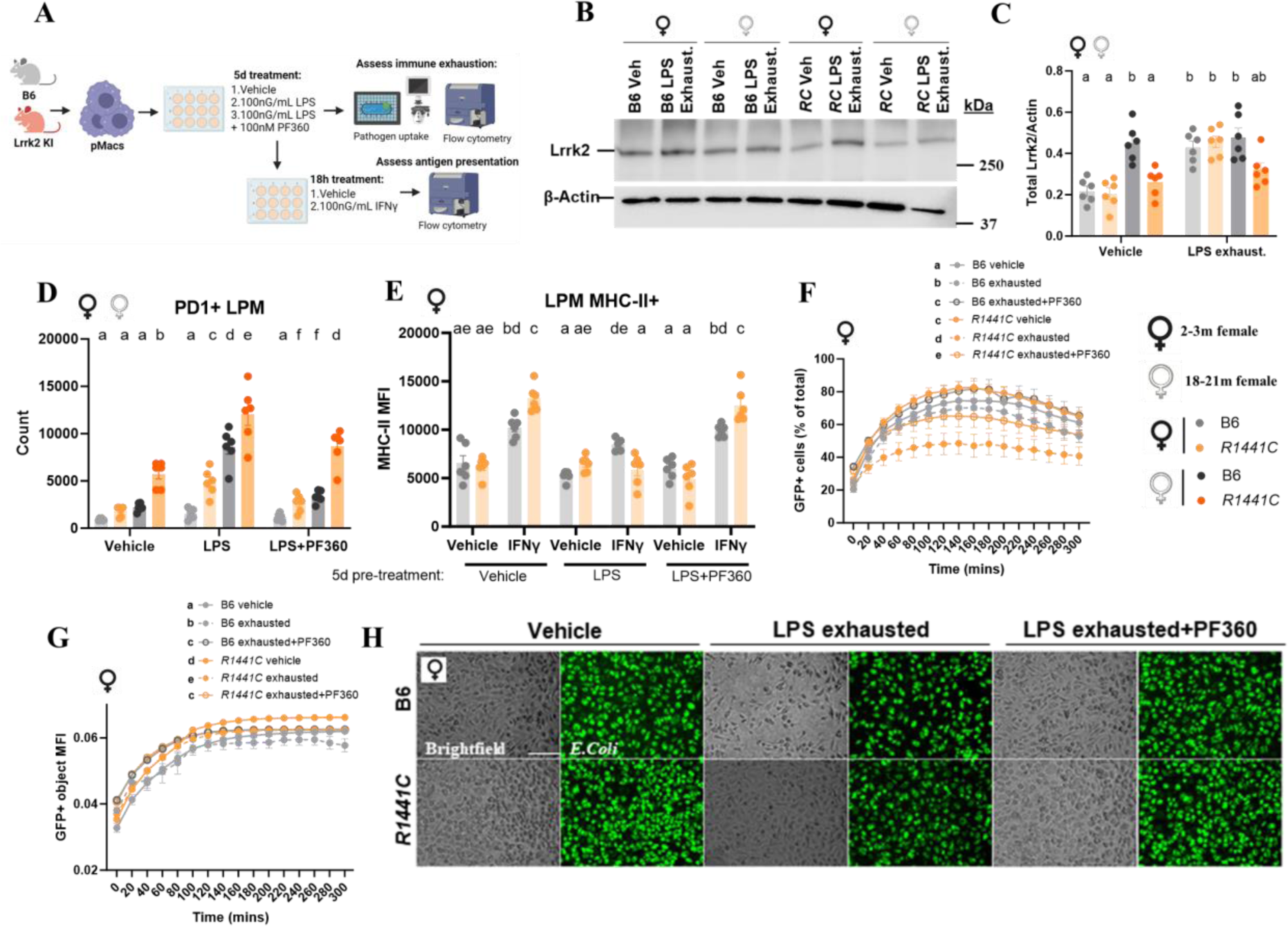
*R1441C* Lrrk2 mutation induces immune cell exhaustion in a kinase-dependent manner. pMacs from 2-3-month and 18-21-month-old female, B6 or *R1441C* mice were incubated with 100ng/mL of LPS for 5 days +/- 100nM PF360. (A) Schematic of study design. (B, C) Lrrk2 protein expression was assessed and normalized to β-Actin and quantified. Representative western blots shown. (D) PDL1+ LPM count was quantified via flow cytometry. (E) After 5-day LPS or vehicle treatment, pMacs were stimulated with 100nG/mL of IFNγ for 18-hours and MHC-II MFI expression was quantified via flow cytometry. Bars represent mean +/- SEM (N = 6-12 mice). Three -way ANOVA, Bonferroni *post hoc*. (F-H) After 5-day LPS or vehicle treatment, pMacs were incubated with 100μg of pHrodo *E. coli*. +/- 100nM PF360. pMacs were imaged every 20 minutes for 5 hours, and GFP+ cells (% of total) and GFP MFI was measured. Scale bars, 100μM. Four-way ANOVA, Bonferroni *post hoc*. Main effects of groups are reported in each graph key. Groups sharing the same letters are not significantly different (p > 0.05) whilst groups displaying different letters are significantly different (p < 0.05).

A number of negative costimulatory markers that are associated with immune cell exhaustion have been identified, including the immune checkpoint molecule, PDL1 (*54*). PDL1 expression was quantified via multi-color flow cytometry in pMacs from young and aged mice to determine if pMacs from aged *R1441C* females were exhausted as hypothesized. Indeed, an increase in PDL1^+^ cells was observed in both vehicle and LPS-exhausted aged *R1441C* LPMs relative to age-matched B6 controls as well as young mice (Fig. 5D). As this was seen in both vehicle and LPS conditions, these exciting findings suggest that pMacs from *R1441C* females are already exhibiting immune cell exhaustion markers, which is then exacerbated by chronic LPS-exhaustion. Interestingly, a significant increase in PDL1^+^ cell count was observed in pMacs from young *R1441C* females in both vehicle and LPS-exhausted conditions relative to age-matched B6 controls (p<0.01) (Fig. 5D). Interestingly, when cells were pushed to exhaustion with Lrrk2 kinase inhibitor co-treatment, a significant reduction in PDL1^+^ cell count was observed (p<0.05). These data suggest that *R1441C* macrophages from female mice may be susceptible to immune cell exhaustion in a Lrrk2-kinase dependent manner.

LPMs from aged *R1441C* male mice, although no significant effect of genotype was observed in vehicle conditions, when pushed to exhaustion with LPS, an increase in PDL1^+^ LPMs relative to age-matched B6 controls, as well as pMacs from young mice, was observed (Fig. S6C). The lack of differences in PDL1^+^ cells between genotypes or ages in male mice suggests there is no effect of the *R1441C* mutation on immune cell exhaustion in naïve animals. However, based on the increased PDL1^+^ cell counts observed in aged *R1441C* males when treated with LPS for 5 days, we conclude that a secondary, chronic inflammatory insult can push *R1441C* pMacs from male mice to acquire immune cell exhaustion like that seen in female mice.

We next sought to determine if LPS-induced exhausted macrophages from young female *R1441C* mice also exhibited the diminished antigen presentation capabilities that we observe in the aged *R1441C* macrophages. After a 5-day treatment with vehicle or LPS +/- PF360, macrophages were treated with IFNγ and MHC-II expression was quantified via multi-color flow cytometry. In LPMs subjected to a 5-day vehicle pre-treatment, the same increased MHC-II expression in *R1441C* macrophages relative to B6 controls was observed (Fig. 5E). As hypothesized, in LPMs pushed to exhaustion with a 5-day LPS pre-treatment, a decreased response to IFNγ was elicited in both genotypes; however, *R1441C* LPMs exhibited decreased MHC-II relative to B6 controls in response to IFNγ, phenocopying what was observed in macrophages with aged mice. Importantly, this could be ameliorated by co-treatment with PF360 during the 5-day pre-treatment, suggesting that the push to immune cell exhaustion and vulnerability of *R1441C* macrophages is dependent on Lrrk2 protein kinase activity. Furthermore, we also observed that pMacs from young *R1441C* females pushed to exhaustion with a 5-day pre-treatment of LPS exhibited a significant reduction in phagocytosis as measured by uptake of pHrodo™ Green *E. coli* relative to B6 controls detected by GFP^+^ cell percentage and GFP MFI which was ameliorated with PF360 co-treatment (p<0.01) (Fig. 5F-H).

Based on the observation that the PDL1^+^ cell count was elevated in LPMs from aged *R1441C* males relative to B6 controls, we hypothesized that a 5-day LPS treatment would induce immune cell exhaustion in pMacs from aged *R1441C* males similar to that seen in female mice. After a 5-day treatment with vehicle or LPS +/- PF30, macrophages from aged male mice were treated with IFNγ and MHC-II expression quantified via multi-color flow cytometry. LPMs from aged males pushed to exhaustion with 5-day LPS pre-treatment, *R1441C* LPMs exhibited decreased MHC-II relative to B6 controls in response to IFNγ, phenocopying what was observed in macrophages from aged female *R1441C* mice (Fig. S6E). Additionally, the same hypophagocytic phenotype observed in aged *R1441C* female pMacs was observed in males once pushed to exhaustion with a 5-day LPS treatment (Fig. S6F-H). Collectively such data support the hypothesis that, with a secondary hit in the form of a chronic, inflammatory stimulus, male *R1441C* mice exhibit increased immune cell exhaustion relative to B6 controls, similar to that seen in naïve *R1441C* female macrophages.

### Transcriptomic profiling reveals age- and sex-specific alterations in critical effector function pathways in *R1441C* macrophages

To explore the age and sexually dimorphic effects of the *R1441C* mutation on macrophage function, RNA was harvested from pMacs post-vehicle or IFNγ treatment and processed for RNA sequencing (RNAseq). A principal component analysis (PCA) was conducted from the resulting dataset which revealed that groups in our dataset primarily cluster based on age and treatment with vehicle or IFNγ treatment (Fig. S7A).

We first sought to identify key cellular pathways altered under different treatment conditions in *R1441C* pMacs relative to B6 controls. In pMacs from young females treated with IFNγ, Gene Ontology (GO) enrichment analysis revealed that upregulated differentially expressed genes (DEGs) in *R1441C* macrophages were enriched in pathways such as regulation of phagocytosis, T cell differentiation, cytokine production and defense response to virus (Fig. S7B, C). In pMacs from aged females treated with IFNγ, GO enrichment analysis showed downregulated DEGs in *R1441C* macrophages were enriched in pathways such as detection of external biotic stimulus, acute inflammatory response, and regulation of MAP kinase activity (Fig. S7D, E). Collectively, such observations support our previous data that shows an effect of *R1441C* Lrrk2 on functional readouts such as increased phagocytosis and cytokine production in response to IFNγ in young females relative to B6 controls, whilst in aged mice, effector functions responsible for defense responses are significantly downregulated relative to B6 controls.

Overall, fewer genes were dysregulated in *R1441C* pMacs from young females treated with vehicle, relative to pMacs treated with IFNγ (Fig. S8A), although these DEGs were enriched in similar pathways (Fig. S8B). Similar observations were made with pMacs treated with vehicle from aged females (Fig. S8C). As Lrrk2 protein increases in response to IFNγ, it follows that DEGs driven by the *R1441C* mutation would have a more significant biological impact in these conditions where Lrrk2 protein expression are increased. Furthermore, given that the majority of phenotypes we observe in pMacs from *R1441C* females are seen in response to an inflammatory insult or immunological challenge, such as IFNγ or *E. Coli,* it is logical that more *R1441C*:B6 DEGs would be observed in these conditions relative to vehicle treatments. Few *R1441C*:B6 DEGs were observed in pMacs from male mice at both ages and in both treatments (Fig. S8D-I). This is unsurprising, given that many phenotypes were not observed in *R1441C* male mice. pMacs from aged *R1441C* only begin to exhibit phenotypes similar to *R1441C* female mice when pushed to immune cell exhaustion and re-challenged with IFNγ. It is therefore likely that to identify key cellular pathways that are altered by the *R1441C* mutation in male mice, a second, chronic immune insult is needed to elicit those changes.

### *R1441C* Lrrk2 mutation alters transcriptional regulation of mitochondrial and oxidative stress genes in macrophages in a sex- and age-dependent manner

Based on the observation that the *R1441C* mutation alters gene expression in pMacs from aged female mice, we next sought to determine the key cellular pathways that this mutation altered in the aging process. When comparing DEGs in pMacs from both B6 and *R1441C* aged mice treated with IFNγ relative to those from young pMacs, there was a large degree of overlap between the DEGs identified in the two genotypes (Fig. 6A-C). However, 2012 genes were identified to be differentially expressed upon aging and novel to *R1441C* pMacs. KEGG pathway enrichment analysis revealed that these 2012 genes unique to aged *R1441C* pMacs were significantly enriched for pathways associated with neurodegenerative diseases, such as Parkinson’s disease, Alzheimer’s disease and Huntington’s disease (p<0.05) (Fig. 6D). Similarly, GO biological pathway enrichment analysis of these DEGs showed a significant enrichment (p<0.05) in pathways related to mitochondria, oxidative stress and metabolism, including mitochondrial respiratory chain complex I assembly, glucose and pyruvate metabolic process and cellular respiration, which are all known key lysosomal and mitochondrial functions. Many DEGs observed in the Parkinson’s Disease KEGG pathway were present in the GO biological pathways associated with mitochondrial function and oxidative stress, including many *Nduf* and *Cox* genes*, Park7, Pink1* and *Prkn* (Fig. 6E). Furthermore, GO and KEGG pathway enrichment analysis of DEGs identified as upregulated in pMacs from aged *R1441C* females relative to aged *R1441C* males showed a significant enrichment (p<0.05) in these same pathways, including Parkinson’s Disease, cellular respiration and ribonucleotide biosynthetic process (Fig. 6F). Given that pMacs from aged *R1441C* females are susceptible to the age-dependent, bi-phasic immunophenotype described here, whilst pMacs from aged *R1441C* males seem to be protected, altered mitochondrial function as a function of age in the females, which is not observed in the males, may underlie this observation.

**Figure 6.**
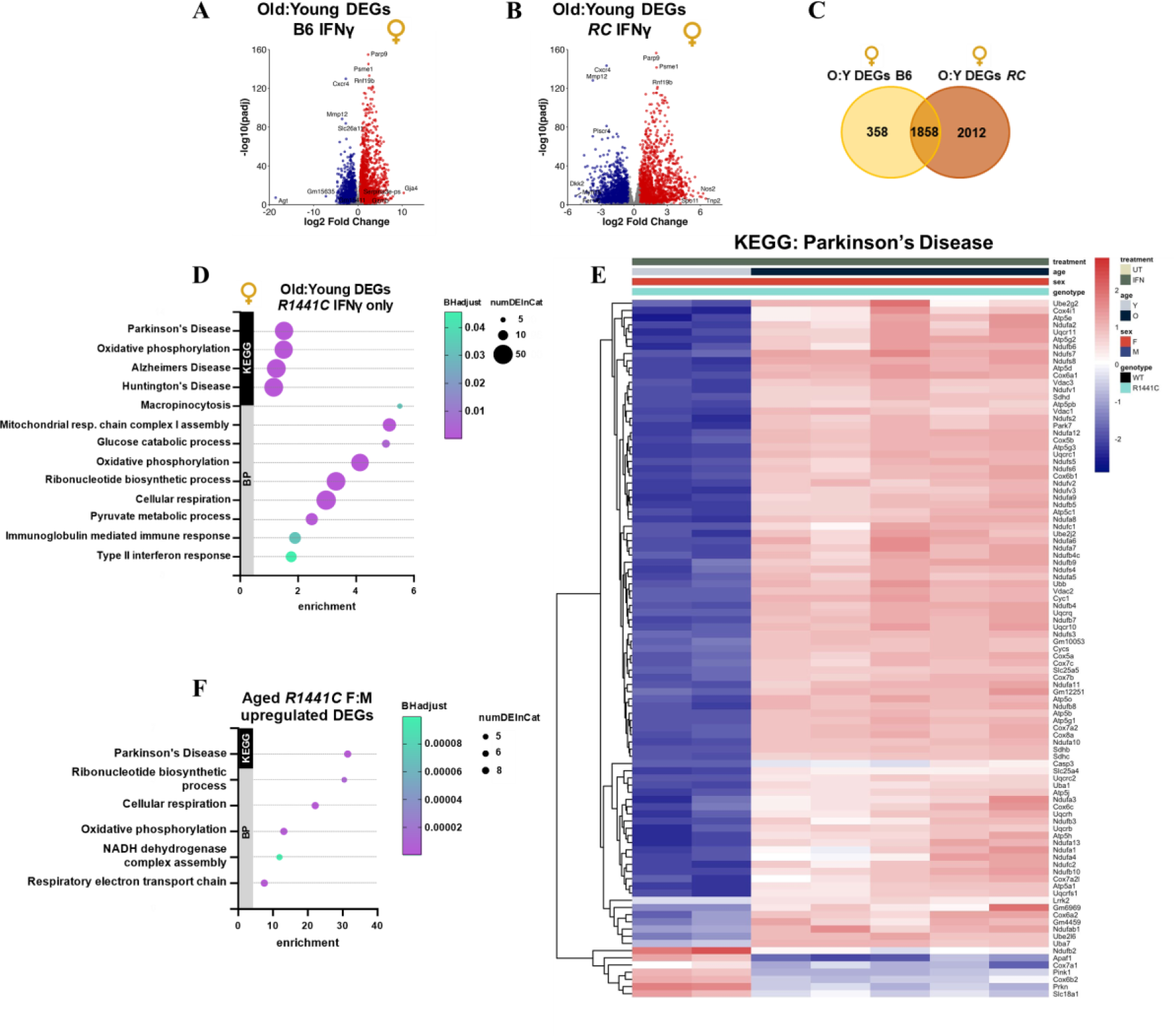
*R1441C* Lrrk2 mutation alters transcriptional regulation of mitochondrial and oxidative stress genes as a function of age in macrophages from mutant females. Transcriptomic analysis from B6 or *R1441C* vehicle and IFNγ-treated pMacs, harvested from 2-3-month or18-21-month-old. (A, B, C). Significant Old:Young DEGs in IFNγ-treated pMacs were counted and compared across B6 and *R1441C.* Volcano plots show genes with log_2_-transformed fold change > 0.5 and an adjusted p-value ≤ 0.05. (D) KEGG pathway and GO BP enrichment analysis identifies pathways in Old:Young DEGs seen only in *R1441C* samples. (E). Heat maps show KEGG: Parkinson’s Disease pathway with significant Old:Young DEGs seen only in *R1441C* samples. (F) KEGG pathway and GO BP enrichment analysis identifies pathways in Aged R1441C F:M upregulated DEGs. pMac-derived RNA was used from 4-6 mice per sex, genotype, treatment and age in transcriptomic studies.

When comparing DEGs in aged pMacs from both B6 and *R1441C* female mice treated with vehicle from, unsurprisingly, different DEGs were identified relative to what was seen in IFNγ conditions (Fig. S9A-D). As opposed to pathways related to mitochondria and oxidative induced by IFNγ conditions, DEGs novel to *R1441C* in vehicle conditions were significantly enriched (p<0.05) in pathways such as wound healing responses, and Wnt, ERK and PI3K signaling. It seems, therefore, that at rest, or under vehicle-treatment conditions, the *R1441C* mutation exerts its effects on aging through pathways crucial for metabolism, cell survival, and cell growth. The Wnt signaling cascade is crucial in macrophages and serves as a regulator of macrophage inflammation-resolving state (*55*). However, during an inflammatory insult, the *R1441C* mutation exerts its effects on aging through pathways crucial for oxidative stress and cellular respiration.

Interestingly, when assessing DEGs in pMacs from aged male mice treated with IFNγ, 540 genes were identified as novel and upregulated in *R1441C* macrophages that were not significantly affected by aging in B6 controls (Fig. S9E-G). Pathway enrichment analysis revealed that these 540 upregulated genes significantly enriched pathways (p<0.05) related to type II interferon production, cytokine production, and NFkB and JNK signaling (Fig. S9H). We also observed that, with the 5-day LPS stimulation, pMacs from aged *R1441C* male mice could be pushed to exhibit phenotypes that paralleled those seen in aged female *R1441C* macrophages. It has previously been reported that sustained and chronic STAT1 signaling, known to be stimulated by cytokine signaling (*56*), is necessary to induce macrophage exhaustion (*48*). Collectively, we interpret our findings that the upregulated DEGs, including driving increased cytokine release in *R1441C* aged male macrophages, may predispose these macrophages to become exhausted upon LPS-stimulation.

### *R1441C*- and *Y1699C-LRRK2* monocyte-derived patient macrophages exhibit hypophagocytosis and increased markers for exhaustion

To assess the effects of PD-associated mutations on immune cell exhaustion in humans carrying *LRRK2* mutations, PBMCs were collected from 2 PD-*Y1699C* and 1 PD*-R1441C* patient, 9 healthy controls, and 3 non-manifesting carriers (NMC) of the *R1441C* pathogenic variant (Fig. 7A). *R1441C* and *Y1699C* variant are located in the GTPase domain of the LRRK2 protein, and both have been shown to perturb LRRK2’s GTPase activity and have been reported to elevate its kinase activity (*15, 23, 24, 57, 58*). In the cohorts available for this study, although healthy controls were balanced for sex (5 male and 4 female), all PD patients and NMCs were male (Table S3).

**Figure 7.**
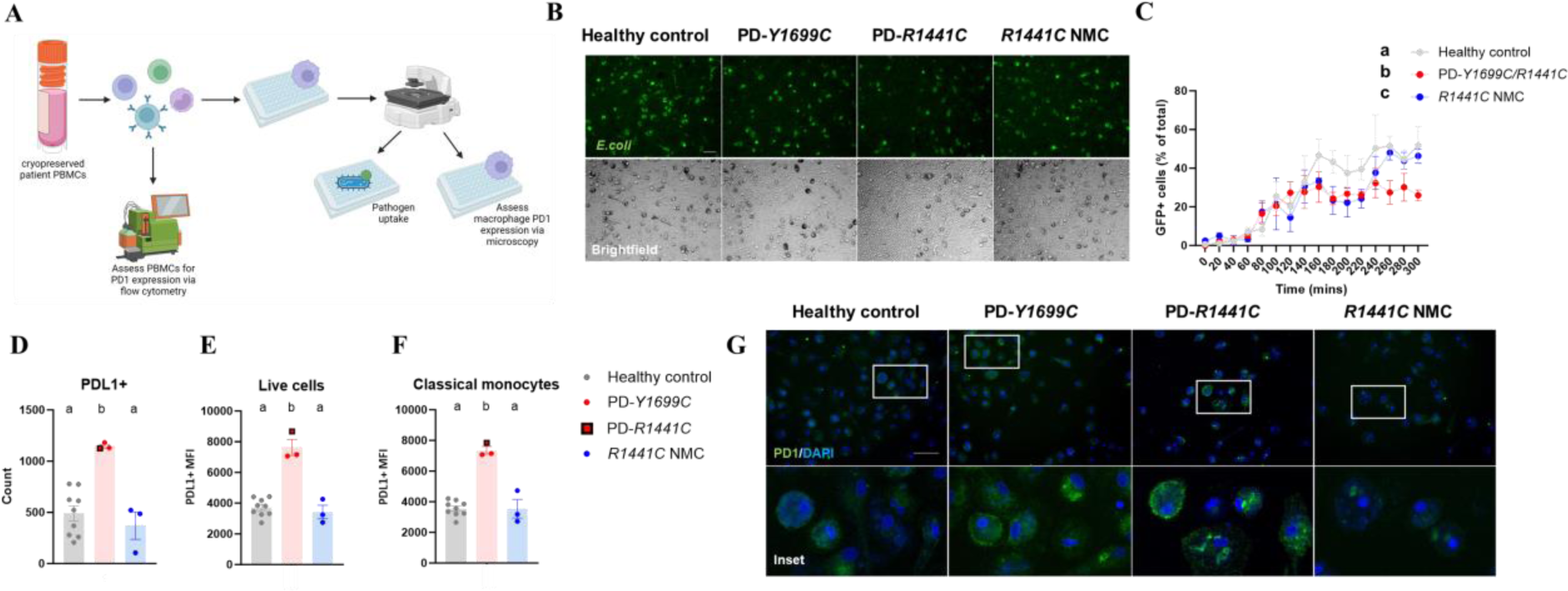
*R1441C* and *Y1699C-LRRK2* monocyte-derived patient macrophages exhibit hypophagocytosis and increased markers for immune cell exhaustion. (A) Schematic of study design. (B, C) Cryopreserved PBMCs were cryorecovered and plated in the presence of hM-CSF to differentiate myeloid cells to macrophages. Macrophages were incubated with 100μg of pHrodo *E. coli* and imaged every 20 minutes for 5 hours, and GFP+ cells (% of total) quantified. . Scale bars, 100μM. Three-way ANOVA, Bonferroni *post hoc*. Main effects of groups are reported in each graph key. Groups sharing the same letters are not significantly different (p > 0.05) whilst groups displaying different letters are significantly different (p < 0.05). (D, E, F) PDL1+ cell count and PDL1 expression was quantified in total PBMCs via flow cytometry. Bars represent mean +/- SEM (N = 3-9 participants). One-way ANOVA, Bonferroni *post hoc*. Groups sharing the same letters are not significantly different (p > 0.05) whilst groups displaying different letters are significantly different (p < 0.05). (G) Monocyte-derived macrophages were stained for PDL1 and microscopy images captured. Scale bars, 100μM.

Pathogen phagocytosis was quantified in monocyte-derived macrophages of these individuals and a hypophagocytic phenotype was observed in monocyte-derived macrophages from PD-*R1441C* and *Y1699C* relative to healthy controls and *R1441C* NMCs, as indicated by a reduced number of GFP^+^ cells (Fig. 7B, C), with no differences in GFP MFI observed between groups (Fig. S10A). Next, to assess whether increased markers of immune cell exhaustion accompanied reduced rates of phagocytosis, PDL1 expression was assessed in peripheral blood mononuclear cells (PBMCs). Consistent with our mouse data, we observed that PBMCs from both PD-*Y1699C* and PD-*R1441C* patients exhibited increased PDL1^+^ cell count and MFI (Fig. 7D, E). When quantifying PDL1 expression in specific immune-cell subsets (Fig. S10B), it was determined that this overall increase in PDL1 expression was driven by expression in classical monocytes, with increased PDL1 expression observed in PD-*Y1699C* and *R1441C* classical monocytes relative to healthy controls and *R1441C* NMCs (Fig. 7F), with no significant differences observed in other cell types (Fig. S10C, D). Similarly, increased PDL1 expression was evident in monocyte-derived macrophages from PD-*Y1699C* and *R1441C* patients relative to healthy controls and *R1441C* NMCs (Fig. 7G). Collectively, the immune cell exhaustion phenotype observed in aged *R1441C* female mice is also associated with an exhaustion phenotype in cells from patients harboring the *LRRK2*-*Y1699C* or *R1441C* mutation.

## DISCUSSION

Considering age is the greatest risk factor for many neurodegenerative diseases, aging, in particular aging of the immune system, is the most underappreciated and understudied contributing factor in the neurodegeneration field. Our novel findings reveal a bi-phasic age- and sex-dependent effect of the *R1441C* mutation in peripheral immune cells, with increased cytokine release, antigen presentation, phagocytosis and lysosomal activity in macrophages from young female *R1441C* mice, with many of these phenotypes ameliorated by Lrrk2 protein kinase inhibition. Our findings further reveal an age-acquired immune cell exhaustion in macrophages from *R1441C* females associated with hypophagocytosis, decreased antigen presentation, and reduced lysosomal pan-cathepsin activation and protein degradation, increased pathogen-mediated DNA breakage, and increased PDL1 expression indicative of immune cell exhaustion. Importantly, this immune cell exhaustion phenotype was also observed in monocytes and macrophages from human subjects with PD and harboring the *R1441C* and the *Y1699C* mutation. Furthermore, it was demonstrated that, although macrophages from male mice did not exhibit this bi-phasic age-dependent phenotype, they could be pushed to exhaustion with chronic LPS stimulation, suggesting that, along with age, a secondary inflammatory insult is required to observe immune cell exhaustion phenotypes in macrophages from *R1441C* males. Finally, it was demonstrated that there is an age-acquired dysregulation in the expression of genes associated with PD risk, mitochondrial function, and oxidative stress in macrophages from female *R1441C* mice.

Consistent with findings in *ex vivo* studies in macrophages, we consistently observed elevated expression of both IL10 and IL6 in circulating plasma from young *R1441C* mice. It has recently been shown that RAW264.7 cells expressing *T1348N*-*LRRK2*, an artificial P-Loop null mutation that disrupts GTP binding (*59, 60*), produce significantly less IL10 relative to wild-type cells in response to LPS and Zymosan (*61*). IL10 is produced by macrophages and is critical in limiting immune-mediated pathology (*62*), and it was therefore suggested by Nazish *et al* that there may be a neuroprotective role of LRRK2 in immune signaling through regulation of IL10 secretion. In addition, we also observed elevations in IL6, which is known to be promptly and transiently produced in response to infections and tissue injuries during host defense through the stimulation of immune reactions (*63*). These observations implicate a protective role of Lrrk2 in young *R1441C* mice. The pro-inflammatory cytokine, TNF, was also increased in circulating plasma of young *R1441C* mice, a compensatory increase that may also contribute to organisms ability to fight infection given that mice lacking TNF exhibit increased susceptibility to infection with increased bacterial load (*64*). In support of the protective role of LRRK2 in immune signaling, LRRK2 has been demonstrated to provide increased pathogen control of *Salmonella typhimurium* (*32, 65*) and *Listeria monocytogenes* (*33*). Consistent with these reports, we observed increased *E. coli* uptake and decreased *E. coli*-mediated DNA damage in macrophages from young *R1441C* female mice, providing further support for a protective role of LRRK2 in immune signaling and defense against pathogens, at least in a young organism. Given that *LRRK2* mutations are associated with not only PD, but also Crohn’s Disease (CD) (*2, 12, 66*) and leprosy (*67*), it is difficult to understand why these mutations would have persisted in the human population. Given the evidence discussed and the data presented here, it is plausible to speculate that these mutations may have persisted due to the immunological advantage they confer to young organisms, including protection against certain infections.

For the first time, we have reported here an age-dependent, bi-phasic effect of LRRK2 on immune cell function, with the *R1441C* mutation inducing myeloid cell immune cell exhaustion in a sex-dependent manner. The concept of immune cell exhaustion was first described as the clonal deletion of virus-specific CD8^+^ T-cells that occurs during high-grade chronic infections, and in T-cells is characterized by the stepwise and progressive loss of T-cell functions (*68*). Regarding exhausted myeloid cells, pathogenic inflammation and immunosuppression, characterized by decreased antigen presentation, hypophagocytosis and altered cytokine release, have been described more recently (*48*). The immune system is composed of numerous cell types with varying states of activation, and cell-to-cell communication during immune responses is crucial for host defense and resolution of inflammation. The observations made here in innate immune cells, therefore, likely have downstream effects on myeloid cell communication with adaptive immune cell types in the periphery. Indeed, alterations in the adaptive immune system have been reported in PD patients. For example, a meta-analysis of 943 cases of PD identified a reduction in circulating CD4^+^ T cells in PD-patients (*69, 70*). More specifically, HLA-DR^+^ T cells and CD45RO^+^ memory T cells have been shown to be increased in people with PD relative to healthy age- and sex-matched controls, while naive CD4^+^ T cells are reduced (*70–72*). CD4^+^FOXP3^+^ Treg cells have increased suppressive activity in patients with PD (*73*). Interestingly, one characteristic of immune cell exhaustion is the dysregulated cross- talk between CD4^+^ and CD8^+^ T-cells due to the suppressive action of Tregs (*74*). Furthermore, Tregs directly inhibit the antigen-presenting function of monocytes and macrophages (*75*). Further research is required to understand the role of cell-to-cell communication in these myeloid cell phenotypes, and if immune cell exhaustion driven by LRRK2 is a contributing factor.

A sex-dependent effect of the *R1441C* mutation was reported here, with females exhibiting an age-acquired immune cell exhaustion phenotype which could only be elicited in macrophages from males upon a secondary, chronic inflammatory insult. Sex-dependent effects of *LRRK2* mutations have previously been reported. A protective effect of the *G2019S* mutation against *S. typhimurium* has been observed in mice, with female mutant mice showing increased survival relative to B6 controls which was not observed in males (*65*). Such observations are also consistent with the sexual dimorphism reported in subjects with PD who also carry a heterozygous p.*G2019S* mutation in *LRRK2*, with female sex being more common amongst carriers than non-carriers (*76–78*). This sexual dimorphism has also been reported for the *R1441C/G/H* mutations (*77*). We observed an *R1441C*-dependent phenotype in macrophages from male mice only when a secondary, chronic inflammatory insult was applied. It is particularly interesting that genes that are significantly upregulated with age in *R1441C* macrophages from male mice are associated with pathways which are heavily implicated in the phenotypes we observed in macrophages from young *R1441C* females. An increase in gene expression in pathways associated with cytokine production is consistent with our previous observation of increased IL6, IL10 and TNF release in aged male pMacs relative to B6 controls. Further research is therefore required to determine the extent to which this sexual dimorphism is driven by altered immune system susceptibility to *LRRK2* mutations in males and females.

We observed alterations in genes enriched in the KEGG Parkinson’s Disease pathway in *R1441C* female macrophages as a function of age, with these genes being mostly associated with mitochondrial respiration and oxidative stress. Not surprisingly, mitochondrial dysfunction has emerged as one of the key mechanisms underlying the pathogenesis of both sporadic PD and familial Parkinsonism (*79, 80*). LRRK2 has been heavily implicated in mitochondrial function and mitophagy, with *LRRK2* mutations causing reduced mitochondria potential and decreased ATP production, mitochondrial DNA damage, mitochondrial fragmentation and altered rates of mitophagy in various cell types and pre-clinical models (*81–84*). But importantly, our findings herein reveal that pathways once thought to be primarily important to neuronal demise (mitochondrial and oxidative stress) are also disrupted in peripheral immune cells by PD-associated mutations in a sex- and age-dependent manner and have important implications for monitoring risk, target engagement, and disease progression of PD and perhaps other NDs by assessing peripheral immune cell types *ex vivo*. It is known that, in T cells, mitochondrial stress induced by chronic antigen stimulation drives immune cell exhaustion (*85*). Further research is therefore required to fully understand the role of mitochondria and mitophagy, and therefore lysosomal function, in the *R1441C*-mediated immune cell exhaustion phenotypes observed here.

GO and KEGG pathway enrichment analysis of DEGs identified as upregulated in pMacs from aged *R1441C* females relative to aged *R1441C* males showed a significant enrichment in pathways related to mitochondrial function and cellular respiration. Interestingly, estrogen increases expression of respiratory complexes, antioxidant molecules, and anti-apoptotic factors that directly impact mitochondrial structure and function, with aging in women being associated with a reduction in estrogen formation and the subsequent development of mitochondrial dysfunction (*86*). Given that pMacs from aged *R1441C* females are susceptible to the age-dependent, bi-phasic immunophenotype described here, whilst pMacs from aged *R1441C* males seem to be protected, altered mitochondrial function as a consequence of aging may underlie this observation. It was also observed that, with chronic-LPS stimulation, pMacs from aged *R1441C* males elicited a phenotype comparable to that seen in pMacs from aged *R1441C* females, suggesting a secondary, chronic inflammatory insult is needed in conjunction with aging in *R1441C* males for immune cell exhaustion to occur. It has previously been reported that upon LPS stimulation, decreased mitochondrial membrane potential decreases and ROS levels increase in macrophages (*87*). Further research is therefore required to decipher if altered mitochondrial function underlies the synergizing effects of aging and chronic inflammation in *R1441C*-induced immune cell exhaustion in macrophages from males.

It has previously been shown that LRRK2 exerts differential effects on inflammatory responses in a tissue-specific manner, and it has been suggested that there is a differential role of LRRK2 in the periphery versus the CNS (*65, 88*). It therefore remains to be determined how the data reported here impacts our understanding of LRRK2 in myeloid cells in both the periphery and the CNS. Indeed, we observed a decrease in MHC-II expression in peripheral myeloid cells of aged *R1441C* females, whilst microglia expressed increased MHC-II in these mice relative to B6 controls, implying a differential effect of LRRK2 in the periphery vs. the CNS. Alterations in type II interferon responses have been observed in both induced pluripotent stem cell (iPSC)-derived microglia and macrophages from patients harboring *G2019S* mutations (*89*), and similar effects of *LRRK2* mutations on lysosomal function have also been reported in patient iPSC-derived microglia and macrophages (*90*). However, it has been demonstrated that expression of *R1441C*-Lrrk2 in peripheral lymphocytes alone is sufficient to induce dopaminergic neuronal cell loss in the midbrains of LPS-treated mice (*39*), suggesting that the periphery plays a pertinent and perhaps primary role in *LRRK2*-PD, and future research into how immune cell exhaustion in the periphery impacts cross- talk with the CNS is needed to inform future immunomodulatory therapeutic interventions for PD.

We successfully demonstrated that, like macrophages from aged *R1441C* mice, myeloid cells from *R1441C* and *Y1699C* patients exhibited hypophagocytosis and increased expression of immune cell exhaustion markers relative to those from HCs and *R1441C*-NMCs. However, it is unclear if the same hyper-responsive phenotype that was observed in young *R1441C* female mice is exhibited in myeloid cells in *R1441C* carriers at a younger age or prior to disease onset. Indeed, we did show that *R1441C*-NMCs did not exhibit these immune cell exhaustion phenotypes, and it is noteworthy that the average age of the 3 *R1441C*-NMCs in this study was 30.5 years (Table S3), whilst the average age of the *R1441C* and *Y1699C*-PD subjects was 42 and 63.5, respectively. Further investigation is therefore required to determine if PBMCs from NMCs, upon follow-up after further aging or disease onset, exhibit immune cell exhaustion. As well, it is interesting to note that, despite the immune cell exhaustion phenotype being observed in female *R1441C* KI mice, all the *R1441C* and *Y1699C*-PD patients recruited here were male (Table S3), and PBMCs from these male patients exhibited similar immune cell exhaustion phenotypes observed in female *R1441C* mice. We do, however, observe these immune cell exhaustion phenotypes in macrophages from aged *R1441C* males upon a secondary, chronic inflammatory insult. Such observations support the hypothesis that complex gene-by-environment interactions combine with an ageing immune system to create the ‘perfect storm’ that enables the development and progression of PD (*5*). Further research is therefore required to determine if sex plays a role in this ‘perfect storm’. Furthermore, further research is required to understand if these phenotypes are exclusive to *LRRK2*-pathogenic variants located in the GTPase domain or if they are seen in other *LRRK2* mutant immune cells, such as the more common *G2019S* mutation, as well as in idiopathic PD or sporadic cases.

LRRK2-targeting therapies, including antisense oligonucleotides (ASOs) and kinase inhibitors, have been suggested as a potential therapeutic for *LRRK2*-PD, with many hypothesizing that targeting the increased LRRK2 levels and kinase activity will be beneficial to CNS-related pathology. Our findings suggest that more research is required to understand the location and optimal timing for the administration of these LRRK2-targeting therapeutics. Specifically, if LRRK2 plays a critical role in pathogen and infection control in a pathogen or tissue-specific manner, it may mean that LRRK2-targeting therapies may need to be limited to carriers of *LRRK2* pathogenic variants and/or may need to be delivered in a CNS compartment-targeted manner as opposed to systemically to avoid deleterious collateral damage to immune function. Indeed, the LRRK2 ASO currently being tested in the clinic is being delivered via intrathecal injection directly into the cisterna magna, thus limiting systemic exposure (NCT03976349). Lastly, perhaps somewhat surprisingly, we found that, while LRRK2 kinase inhibition could rescue some of the effector functions of young mutant- LRRK2 mice, it had no beneficial effects on the immune-cell exhaustion phenotypes observed here in macrophages from aged mice; however, it did prove to be protective in macrophages from young mice to prevent the development of an immune-cell exhaustive phenotype. Such observations suggest that LRRK2- targeting therapies would elicit the most beneficial effects during the prodromal phases of disease, and further research on PBMCs from prodromal patients with pre-motor features at risk for development of motor PD due to a family history will be required to investigate this possibility and inform future clinical trials.

Our study is not without limitations, and further steps are needed to translate our findings into clinical samples further. Firstly, although we provide evidence of immune cell exhaustion in patient immune cells, this was carried out in a small population due to the availability of cells to us at the time. It is hard to conclude here whether *R1441C*-NMC did not exhibit immune cell exhaustion phenotypes due to the fact that these phenotypes only present themselves upon disease onset, or, whether immune cell exhaustion phenotypes are not present amongst all *R1441C* carriers. Furthermore, due to the fact that penetrance of these mutations is low, it may be that the NMCs recruited for this study will not develop PD and this explains their lack of immune cell exhaustion phenotypes. Larger, longitudinal studies are therefore required in order to decipher between these possibilities. Secondly, due to the lack of a longitudinal study design and limited sample numbers, it is hard to conclude whether age or chronic immune insult are the primary driver of immune cell exhaustion in our patient samples, and whether immune cell exhaustion is a key driver of PD onset and/or progression. Thirdly, although the *G2019S*-*LRRK2* mutation is the largest contributor to familial forms of PD, and LRRK2 has been implicated in sporadic forms of the disease, the *R1441C* mutation is comparably rarer, with prevalence estimates between 0.03-2% (*91*). The focus on *R1441C*, therefore, may be limited, and further studies are required to determine if these phenotypes are mutation-specific and if they are seen in sporadic and idiopathic-PD cases. Lastly, although increased PDL1 expression and hypophagocytosis were observed in patient samples, there are other effector functions altered due to immune cell exhaustion, and in other cell types, such as cytokine release, T cell proliferation and differentiation. Further studies in clinical and pre-clinical samples are therefore required to understand the effects of LRRK2 mutations on other immune cell effector functions and how immune cell exhaustion may be implicated in this.

## MATERIALS AND METHODS

### Study design

The primary objective of this research was to investigate the effects of the *R1441C LRRK2* pathogenic variant on myeloid cell effector function in a sex- and age-dependent manner. We hypothesized that a biphasic, age-dependent effect of the *R1441C* pathogenic variant would be observed, affecting antigen presentation, cytokine release, phagocytosis and lysosomal function in macrophages. To explore our hypothesis, we used *R1441C* KI mice and B6 controls and assessed myeloid cell effector function both *ex vivo* and *in vivo,* in males and females at 2-3 months and 18-21 months. All studies were performed by investigators blinded to genotype and treatment groups at the University of Florida, USA. Power analyses based on the labs pilot and published data indicated that n = 8-12 per group for flow-cytometry and cytokine release assays, and n = 4-6 for imaging based immunophenotyping and immunoblotting would be needed to yield 80% power with a α set to 0.05. Exclusion criteria for experiments involving mice included mice who reached endpoint before time of experiment (due to health-related issues), mice who were found to have tumors whilst collecting tissue, and datapoints that fell more than 3SD +/- of the average of that group. Multiple cohorts of mice (each cohort consisting of 6-12 mice per group depending on the assay) were used to generate this data set. As 2 sexes, 2 genotypes and 2 ages were used to generate this data, each cohort consisted of 8 groups, equating to 48-96 mice. Each cohort was separated into batches and experiments carried out, and data acquired and analyzed for each batch, with data from batches merged post-analysis to form the overall data set for that cohort. As well, entirely independent cohorts of mice were used to produce data using orthogonal assays (e.g. 1 cohort used for flow cytometry, 1 cohort used for microscopy and western blotting analysis) to ensure the phenotypes we were observing were not assay- or cohort-dependent. By generating this data across batches of animals and using multiple orthogonal assays to confirm phenotypes in separate cohorts of mice, we ensure that our data is robust and reproducible. A schematic of this study design is shown in Fig.S11.

The study used human PBMCs received from collaborators at University Hospital Schleswig-Holstein, University of Luebeck, Germany and the University Hospital Eppendorf, Hamburg, Germany. Samples were received at the University of Florida and investigators were unblinded after completion of analysis. Deidentified data provided included sex, age at collection, age of diagnosis, clinical diagnosis and genetic status.

### Animals

*R1441C* knock-in (Strain #009346; B6.Cg-Lrrk2tm1.1Shn/J) mice have previously been characterized (*92*) and were maintained in the McKnight Brain Institute vivarium (University of Florida) at 22°C at 60-70% humidity with a 12-hour light/dark cycle and *ad libitum* access to standard rodent chow and water. C57BL/6 (Strain #000664) controls were used for all studies. All animal procedures were approved by the University of Florida Institutional Animal Care and Use Committee and were in accordance with the National Institute of Health *Guide for the Care and Use of Laboratory Animals* (NIH Publications No. 80-23) revised 1996. Female and male mice were aged to 2-3 or 18-21-months-old and sacrificed via cervical dislocation or rapid decapitation.

### Human subjects

This study was reviewed and approved by the University of Lubeck Institutional Review Board (AZ16- 039). Participants provided written informed consent to participate. During recruitment, a family history was used to assess history of disease and patients were screened for *LRRK2* variants. See Table S3 for patient demographics.

### Statistics and data analysis

All individual level data can be found in data file S1 and is also available on Zenodo (10.5281/zenodo.11099669). Data and statistical analyses were performed using IBM SPSS statistics 27 or GraphPad Prism 9. For assessing differences between groups, data were analyzed by either 1-way or 2-way analysis of variance (ANOVA), or by t-test. In instances when data did not fit parametric assumptions, Kruskal-Wallis non-parametric ANOVA was used. *Post hoc* tests following ANOVAs were conducted using Tukey HSD or Bonferroni correction. Two-tailed levels of significance were used and p < 0.05 was considered statistically significant. Graphs are depicted by means +/- standard error of the mean (SEM).

## List of Supplementary Materials

### Materials and Methods

#### Supplementary figures

Figure S1. Characterization of ex vivo pMac cultures.

Figure S2. R1441C mutation leads to age dependent biphasic alteration in antigen presentation.

Figure S3. The R1441C Lrrk2 mutation causes alterations of monocyte population and microglia activation in aged female mice.

Figure S4. R1441C mutation leads to age-dependent biphasic alteration in lysosomal activity.

Figure S5. R1441C Lrrk2 mutation alters pathogen uptake and pathogen-mediated DNA damage in age- and sex-dependent manner.

Figure S6. R1441C Lrrk2 mutation induces immune cell exhaustion in a kinase-dependent manner.

Figure S7. Transcriptomic profiling reveals alterations in critical effector function pathways in R1441C macrophages in age- and sex- dependent manner.

Figure S8. R1441C Lrrk2 mutation alters biological processes in macrophages in a sex- and age-dependent manner.

Figure S9. R1441C Lrrk2 mutation alters transcriptional changes associated with age in sex dependent manner.

Figure S10. R1441C- and Y1699C-LRRK2 monocyte derived macrophages exhibit hypo phagocytosis and increased markers for immune cell exhaustion.

Figure S11. Schematic of study design with mice.

#### Supplementary tables

Table S1. Flow cytometry marker antibody panel (mouse)

Table S2. Antibodies for immunoblotting

Table S3. Patient demographics

Table S4. Flow cytometry marker antibody panel (human)

## Acknowledgments

We thank members of the Tansey lab for useful discussions and edits of the manuscript. We thank the UF Interdisciplinary Center for Biotechnology Research (UF | ICBR) for use of flow cytometry and sequencing facilities and advice. Partial funding for this work was derived from awards from the Bright Focus Fellowship A2021017F (RLW), NIH/NINDS grant RF1NS128800 (MGT), NIH NIA 1RF1AG057247, and the joint efforts of The Michael J. Fox Foundation for Parkinson’s Research (MJFF) and the Aligning Science Across Parkinson’s (ASAP) initiative. MJFF administers the grant ASAP-020527 on behalf of ASAP and itself. For the purpose of open access, the author has applied a CC-BY public copyright license to the Author Accepted Manuscript (AAM) version arising from this submission. N.B. was supported by the DFG (BR4328.2-1, GRK1957), the Michael J Fox Foundation, the Collaborative Center for X-linked Dystonia-Parkinsonism and the EU Joint Programme - Neurodegenerative Disease Research (JPND)

## Author contributions

Conceptualization: RLW, MGT

Methodology: RLW, HAS, KMF, SS, NB, SZ, TU, CK, ES

Investigation: RLW, KMF

Visualization: RLW, MGT

Funding acquisition: RLW, MGT

Project administration: MGT Supervision: MGT

Writing – original draft: RLW, HAS, KMF, SS, NB, SZ, TU, CK, ES, MGT

All authors reviewed and approved the final manuscript.

## Data and materials availability

All data are available in the main text or supplementary files, including links to data repositories and individual level data in Data File S1. Correspondence and requests for materials should be addressed to MGT.

