## Supplementary material for "The *R1441C-LRRK2* mutation induces myeloid immune cell exhaustion in an age- and sex-dependent manner": sup materials and figures

**Supplementary materials**

**Materials and Methods**

***Harvesting and culturing of peritoneal macrophages and ex-vivo stimulation***

Peritoneal macrophages were harvested from mice that had received a 1 ml intraperitoneal administration of 3% Brewer thioglycolate broth 72 hours prior collection. Mice also received Buprenorphine Sustained-Release every 48 hours for pain relief. Mice were sacrificed via cervical dislocation and abdomen sprayed with 70% Ethanol. The skin of the abdomen was split along the midline, taking care to avoid puncturing or cutting the abdominal cavity. 10 mL of cold RPMI media (Gibco; 11875119) was injected into the peritoneal cavity using a 27G needle. After gentle massaging of the peritoneal cavity, as much fluid was withdrawn as possible from the peritoneal cavity using a 25G needle and 10 mL syringe. Aspirated fluid was passed through a 70uM nylon filter onto 50mL falcon and pre-wet with 5mL of HBSS-/-. Filters were then washed twice with 5mL of HBSS-/- and then tubes spun at 400*x g* for 5 minutes at 4°C. Supernatant was aspirated and pellet resuspended in 3 mL pre-warmed growth media (RPMI, 10% FBS, 1% Pen-Strep). Cells were counted and viability recorded using trypan-blue exclusion on an automated cell-counter (Countess™; Thermo). Volume growth media was adjusted so that cells were plated at 5 x $10^5$/mL in 6-, 24- or 96-well plates depending on the intended assay. Cells were incubated at 37°C, 5% CO2 for a minimum of 2-to-3 hours to allow macrophages to adhere. Wells were washed twice with sterile PBS to remove non-adherent cells and new, pre-warmed growth media added. For cells requiring *ex vivo* stimulation, 100U of IFNγ (R&D) or vehicle ($H_2O$) was added for 18 hours. For co-treatments, a final concentration of 40 nM Bafilomycin A1 (Sigma) and 100 nM PF360 (PF-06685360; MedChem) was used. Protocol available at dx.doi.org/10.17504/protocols.io.j8nlkoyoxv5r/v1.

***Flow cytometry***

1-hour prior to collection, BMV109 Pan Cathepsin probe (Vergent Bioscience) and DQ red BSA (Invitrogen) were added to each well at a final concentration of 1µM and 10µg/mL, respectively, and cells were incubated at 37°C for 1-hour. Cells were then washed 3 times in sterile PBS, harvested, and transferred to a v-bottom 96-well plate (Sigma, CLS3896-48EA) and centrifuged at 300*x g* for 5 minutes at 4°C. Cells were resuspended in 50 µL of PBS containing diluted fluorophore-conjugated antibodies (see Table S1) and incubated in the dark at 4°C for 20 minutes. Cells were centrifuged at 300*x g* for 5 minutes at 4°C and washed in PBS x 2. Cells were fixed in 50 µL of 1% paraformaldehyde (PFA) at 4°C in the dark for 30 minutes. Cells were centrifuged at 300*x g* for 5 minutes and resuspended in 200 µL FACs buffer (PBS, 0.5 mM EDTA, 0.01% sodium azide). Cells were taken for flow cytometry on a Macs Quant Analyzer (Miltenyi) or BD LSRFortessa™ Cell Analyzer. A minimum of 100,000 events were captured per sample and data were analyzed using FlowJo version 10.6.2 software (BD Biosciences; RRID:SCR_008520). When validating multi-color flow cytometry panels and antibodies, fluorescence minus one controls (FMOCs) were used to set gates and isotype controls were used to ensure antibody-specific binding. Compensation was achieved using SPHERO Supra Rainbow beads (Spherotech, Inc.) by setting laser voltages to achieve fluorescent intensity measurements in each channel equal to those in previous experiments. Protocol available at dx.doi.org/10.17504/protocols.io.rm7vzx9x4gx1/v1.

***$Ea_{(52–68)}$ uptake assays***

MHC-II Ea chain (Ea) (52–68) peptide (Anaspec) was reconstituted in sterile distilled $H_2O$ to a final concentrate of 1mg/mL. Once peritoneal macrophages had adhered to plates, 5µg per well was added. Cells were incubated for 18-hours and taken forward for flow cytometry. Protocol available at dx.doi.org/10.17504/protocols.io.14egn3r3pl5d/v1.

***Cytokine release measurements via MesoScale Discovery electrochemiluminescence***

V-PLEX mouse pro-inflammatory panel 1 kit (MSD; K15048D) was used to quantify cytokines in conditioned media from pMacs. Media was diluted 1:1 with MSD kit diluent and incubated at room temperature in the provided MSD plate with capture antibodies for 2 hours as per manufacturer's instructions. Plates were then washed x 3 with PBS with 0.05% Tween-20 and detection antibodies conjugated with electrochemiluminescent labels were added and incubated at room temperature for another 2 hours. After 3 x washes with PBS containing 0.05% Tween-20, MSD read buffer was diluted to 2x and added, and the plates were loaded into the QuickPlex MSD instrument for quantification. Results were normalized to total live cell counts as measured via flow cytometry.

***Intracellular and extracellular MHC-II immunostaining and microscopy***

Methods for intracellular and extracellular MHC-II immunostaining were adapted from previous reports (*42; 10.1016/j.jim.2015.04.023.*). Cells were washed 3x with DPBS+/+ and incubated with 25 ug/mL of APC-MHC-II (Biolegend) in DPBS+/+ containing FcR blocking reagent for 30 minutes at room temperature. Cells were then washed 3x with DPBS+/+ and fixed by incubation in 4% PFA for 10 minutes at room temperature and then washed 3x with DPBS+/+. Fixed cells were permeabilized with permeabilization buffer (eBiosciences) on ice for 15 minutes. 25 ug/mL of PE-610-MHC-II (Biolegend) was spiked in and cells were incubated for 30 minutes at room temperature. Cells were then washed 3 x in DPBS +/+ and incubated in 1 µg/ml DAPI (Life Technologies) for 10 minutes at room temperature in DPBS+/+. Cells were imaged using an EVOS™ M7000 (Invitrogen) at 20 x magnification. Image analysis was performed using Cellprofiler 4.2.5 (RRID:SCR_007358). Using the 'IdentifyPrimaryObject' module in Cellprofiler, icMHC and exMHC were identified in their respective channels and MFI was quantified and Ex:IcMHCII calculated from these quantified MFI values.

***DQ-BSA and BMV109 microscopy***

BMV109 Pan Cathepsin probe (Vergent Bioscience) and DQ red BSA (Invitrogen) were added to each well at a final concentration of 1µM and 10µG/mL, respectively, and cells were incubated at 37°C for 1-hour. Cells were washed 3 x DPBS +/+ and fixed for 10 minutes at room temperature in Invitrogen™ eBioscience™ Intracellular Fixation buffer. Cells were washed 3 x DPBS+/+ and incubated in 1 µg/ml DAPI (Life Technologies) for 10 minutes at room temperature in DPBS+/+. Cells were imaged using an

EVOS™ M7000 (Invitrogen) at 20 x magnification. Image analysis was performed using Cellprofiler 4.2.5. Protocol available at dx.doi.org/10.17504/protocols.io.261gedk5ov47/v1.

***Immunoblotting***

Media was aspirated and cells washed in PBS and lysed in RIPA buffer (50 mM Tris pH 8, 150 mM NaCl, 1% NP-40, 0.5% NaDeoxycholate, 0.1% SDS). Cell lysates were then centrifuged at 10,000*x g* for 10 mins at 4°C. 6X Laemmli sample buffer added (12% SDS, 30% β-mercaptoethanol, 60% Glycerol, 0.012% Bromophenol blue and 375 mM Tris pH 6.8) and samples were reduced and denatured at 95°C for 5 minutes. Samples were loaded into 4-20% Criterion Tris-HCl polyacrylamide gels (BioRad, #3450033) alongside Precision plus protein dual-color ladder (Biorad) to determine target protein molecular weight. Electrophoresis was performed at 100 V for ~60 minutes and proteins transferred to a polyvinylidene difluoride (PVDF) membrane using a Trans-Blot Turbo Transfer System (BioRad) which utilizes Trans-Blot Turbo Midi PVDF transfer packs (BioRad; #1704157) in accordance to manufacturer's instructions. Prior to blocking, total protein was measured using Revert total protein stain (Licor) and imaged on the Odyssey FC imaging system (Licor). Membranes were then blocked in 5% non-fat milk in TBS/0.1% Tween-20 (TBS-T) for 1 hour at room temperature and subsequently incubated with primary antibody (see Table S2) in blocking solution overnight at 4°C. Membranes were washed with TBS-T (3 x 5 minutes) and incubated in horseradish peroxidase (HRP)-conjugated secondary antibody (1:5000) (BioRad) in blocking solution for 1 hour. Membranes were washed in TBS-T (3x 5 minutes) and developed using Super signal west femto/pico (Thermo). Membranes were imaged using the Odyssey FC imaging system and quantified using Image Studio Lite Version 5.2 (Licor; RRID:SCR_013715).

***pH-Rodo pathogens to monitor phagocytosis***

pHrodo™ Green *E. coli* BioParticles™ were resuspended in Invitrogen™ Live Cell Imaging Solution to a final concentration of 1mg/mL and sonicated in bath sonicator for 10 minutes. 100 µL of *E. coli* BioParticles solution was added to each well (96-well plate). Cells were placed in a live imaging chamber (37°C, 5%

carbon dioxide, and 95% humidity) of an EVOS™ M7000 (Invitrogen) at 20 x magnification and 4 images per well captured every 20 minutes for 5 hours. Brightfield images were used for automated-focus throughout the time course and used for normalization of GFP *E. coli* images. Protocol available at dx.doi.org/10.17504/protocols.io.q26g7pn18gwz/v1.

#### *In Situ Apoptosis Detection*

Click-iT™ Plus TUNEL Assay Kits for In Situ Apoptosis Detection (Invitrogen, #C10617) was used to detect DNA breaks in pMacs as per manufacturer instructions. pMacs were harvested and transferred to a 96-well v-bottom plate. pMacs were stained with live/dead dye and antibodies as previously described. pMacs were fixed in 4% paraformaldehyde for 15 minutes at room temperature. Samples were centrifuged at 300*x g* at 4°C, supernatant removed and samples resuspended in permeabilization reagent (0.25% Triton™ X-100 in PBS) for 20 minutes at room temperature. Samples were centrifuged at 300*x g* at 4°C and TdT reaction buffer added and incubated at 37°C for 10 minutes. Samples were centrifuged at 300*x g* at 4°C, and TdT reaction mixture containing EdUTP and TdT enzyme added and cells were incubated for 60 minutes at 37°C. Cells were washed 2 x with 3% BSA in PBS. Click-iT™ Plus TUNEL reaction cocktail was added to each sample and incubated for 30 minutes at 37°C. Cells were taken for flow cytometry on a Macs Quant Analyzer (Miltenyi). A minimum of 100,000 events were captured per sample and data were analyzed using FlowJo version 10.6.2 software (BD Biosciences). When validating Click-iT™ Plus TUNEL Assay Kit for flow cytometry, positive controls were used using DNase I to induce DNA strand breaks and unstained cells for negative controls.

#### *Generation of exhausted peritoneal macrophages*

Exhausted macrophages were generated as previously described (*93*). pMacs were harvested and plated at a final concentration of 5 x $10^5$/mL in 6-, 24- or 96-well plates depending on the intended assay. After 2-hours, cells were washed 2 x in sterile PBS to remove non-adherent cells and growth media containing 100ng/mL of lipopolysaccharide (LPS) was added and cells incubated at 37°C for 5 days, with a full media change on day 3 to replace nutrients. After 5 days, cells were washed 2 x in sterile PBS and either harvested for flow cytometry or for antigen presentation assays.

***Patient PBMC collection and cryopreservation***

8 mL of blood was collected from healthy volunteers per BD Vacutainer CPT Cell Preparation Tube with Sodium Citrate (BD Biosciences, 362761). CPT tubes were inverted 8-10 times and centrifuged at room temperature at 1500*x g* for 20 minutes with no brake at room temperature. The PBMC enriched layer was transferred to a new 50-mL conical tube and MACS buffer (PBS, 0.5% bovine serum albumin, 20 mM EDTA, pH 7.2) was added to a final volume of 50 mL, followed by centrifugation at 1800*x g* for 10 minutes at room temperature. Following removal of the supernatant, PBMCs were resuspended in 10 mL MACS buffer and counted using a hemocytometer with Trypan blue at a 1:20 dilution. PBMCs were centrifuged for 10 min at 300*x g* at 4°C. Supernatant was aspirated and cell pellets were gently resuspended in cryopreservation media (RPMI 1640, with 20% FBS and 10% DMSO) at a final concentration of 1 x $7^7$ cells/mL in cryovials (Simport, T311-2). Cryovials were placed in a room-temperature Mr. Frosty freezing container with isopropanol as per manufacturer's instructions and stored at -80°C overnight. Cryovials were removed from freezing containers and immediately placed into liquid nitrogen for long-term storage. All cryovials were shipped on dry ice and temperatures monitored to ensure the integrity of samples were retained.

***Patient PBMC cryorecovery***

Patient PBMCs were cryorecovered as previously described (*94*). PBMCs were retrieved from liquid nitrogen, thawed at 37°C, slowly added to 37°C filter sterilized complete culture media (RPMI 1640 media, 10% heat-inactivated FBS, 1mM Penicillin-Streptomycin) and pelleted via centrifugation at 300*x g* for 10 min at room temperature. The supernatant was removed and cells were resuspended in 37°C complete culture media for cell counting using Trypan blue.

***Patient PBMC differentiation to macrophages***

PBMCs were plated at a final concentration of 3 x $10^6$/mL in a poly-D-lysine coated, 96-well plate, resulting in 3x$10^5$ cells per well. PBMCs were incubated at 37°C for 2-hours to allow adherent cells to adhere. Cells were washed 2 x sterile PBS to wash off non-adherent cells, and incubated in growth media containing 20ng/mL of human macrophage-colony stimulating factor (hM-CSF) for 5 days, with a full media change on day 3 to replace nutrients. Protocol available at dx.doi.org/10.17504/protocols.io.6qpvr3w83vmk/v1.

***Patient PBMC flow cytometry***

1 x $10^6$ PBMCs were taken for flow cytometry and transferred to a v-bottom 96-well plate (Sigma, CLS3896-48EA) and centrifuged at 300*x g* for 5 minutes at 4°C. Cells were resuspended in 50 µL of PBS containing diluted fluorophore-conjugated antibodies (see Table S4) and incubated in the dark at 4°C for 20 minutes. Cells were centrifuged at 300*x g* for 5 minutes at 4°C and washed in PBS x 2. Cells were fixed in 50 µL of 1% paraformaldehyde (PFA) at 4°C in the dark for 30 minutes. Cells were centrifuged at 300*x g* for 5 minutes and resuspended in 200 µL FACs buffer (PBS, 0.5 mM EDTA, 0.1% sodium azide). Cells were taken for flow cytometry on a MACS Quant Analyzer (Miltenyi). A minimum of 100,000 events were captured per sample and data were analyzed using FlowJo version 10.6.2 software (BD Biosciences). When validating flow cytometry panels and antibodies, fluorescence minus one controls (FMOCs) were used to set gates and isotype controls were used to ensure antibody-specific binding Protocol available at dx.doi.org/10.17504/protocols.io.n92ldm688l5b/v1.

***RNA extraction from pMacs, library prep and sequencing***

RNA from approximately 2-4 x $10^6$ cells was isolated. RNeasy mini kit (Qiagen, # 74104) was used according to manufacturer's instructions. Briefly, 10 µL of β-mercaptoethanol was added to every 1 mL of RLT buffer. 350 µL was added to each well and cells homogenized manually with a mini cell scraper. Cell lysate was transferred to an RNAase free Eppendorf and an equal volume of 70% ethanol was added to each sample. Samples were loaded into supplier columns and centrifuged at 11,000 x *g* for 30 seconds and flow through discarded. 350 µL RW1 buffer was added to each column and centrifuged at 11,000 x *g* for 15 seconds and flow through discarded. RW1 buffer was added and columns centrifuged at 11,000 x *g* for 15 seconds. RPE buffer was added to columns, centrifuged for 11,000 x *g* for 30 seconds, repeated, and 30 µL RNase-free water was added and RNA was eluted. RNA concentration was quantified and 260/230 and 260/280 ratios recorded using a spectrophotometer. RIN values were assessed to ensure RNA integrity using Agilent RNA 6000 Nano Kit (Agilent) and Agilent 2100 Bioanalyzer. RNA was sent to the Interdisciplinary Center for Biotechnology Research (ICBR) core at the University of Florida, where library preparation was performed following the NEBNext Ultra II Directional RNA Library Prep Kit for Illumina manual and 2x150bp paired-end sequencing completed on an Illumina NovaSeq 6000 (Illumina).

***FASTQ alignment, gene counts, and differential expression analysis***

FASTQ files were aligned against the mouse genome (GRCm39) and GRCm39.107 annotation using STAR (RRID:SCR_004463) (*95*) to generate BAM files. Gene counts were generated from BAM files using Rsamtools (Bioconductor package; RRID:SCR_024257) and the summarizeOverlaps function with the GenomicAlignments package v1.36.0 (RRID:SCR_024236) (*96*). Differential gene expression analysis was performed with DESeq2 package v1.40.2 (RRID:SCR_015687) using the "DESeq" function with default settings (*97*) which fits a generalized linear model for each gene. Subsequent Wald test P-values are adjusted for multiple comparisons using the Benjamini–Hochberg method (adjusted P-value). Pair-wise changes in gene expression between groups were used to identify DEGs. DEGs will be defined as an absolute log2 fold change ≥0.5 and an adjusted P-value ≤0.05.

***Functional annotation of DEGs***

Gene Ontology (GO) (RRID:SCR_002811) enrichment analysis was performed with goseq (RRID:SCR_017052) (*98*) to identify enrichment in gene ontology categories and KEGG (RRID:SCR_012773) pathways. For DEGs, up- and down-regulated gene lists were analyzed separately. Over-represented P-values were adjusted for multiple comparisons using the Benjamini–Hochberg (BH) adjustments for controlling false-discovery rates. An enrichment score was calculated using an observed-over-expected ratio for each gene list. GO-BP categories (RRID:SCR_002811) and KEGG pathways were plotted if their BH adjusted FDR reached >= 0.05 and the number of DEGs within each category/pathway was greater than 4. To eliminate larger, broad GO-BP parent categories, categories with more than 250 total genes were not plotted. To eliminate redundancy, pathways with significant overlap regarding genes and semantics were merged.

***Analysis of overlapping genes***

Lists of DEGs were compared using VennDiagram v1.7.3 (RRID:SCR_002414) and ggvenn v0.1.10 (https://cran.r-project.org/package=ggvenn) in R to find overlapping and exclusive genes for each comparison. For other graphs, analyses were conducted in R v4.3.1 (RRID:SCR_001905) using the following packages not already mentioned: pheatmap v1.0.12 (RRID:SCR_016418) to generate heatmaps; PCAtools v2.12.0 for PCA plots (https://www.bioconductor.org/packages/release/bioc/html/PCAtools.html); ggplot2 v3.4.3 (RRID:SCR_014601), using ggrepel v0.9.3 (RRID:SCR_017393) and scales v1.2.1 packages (RRID:SCR_019295) for most other plots.

### Supplementary figures

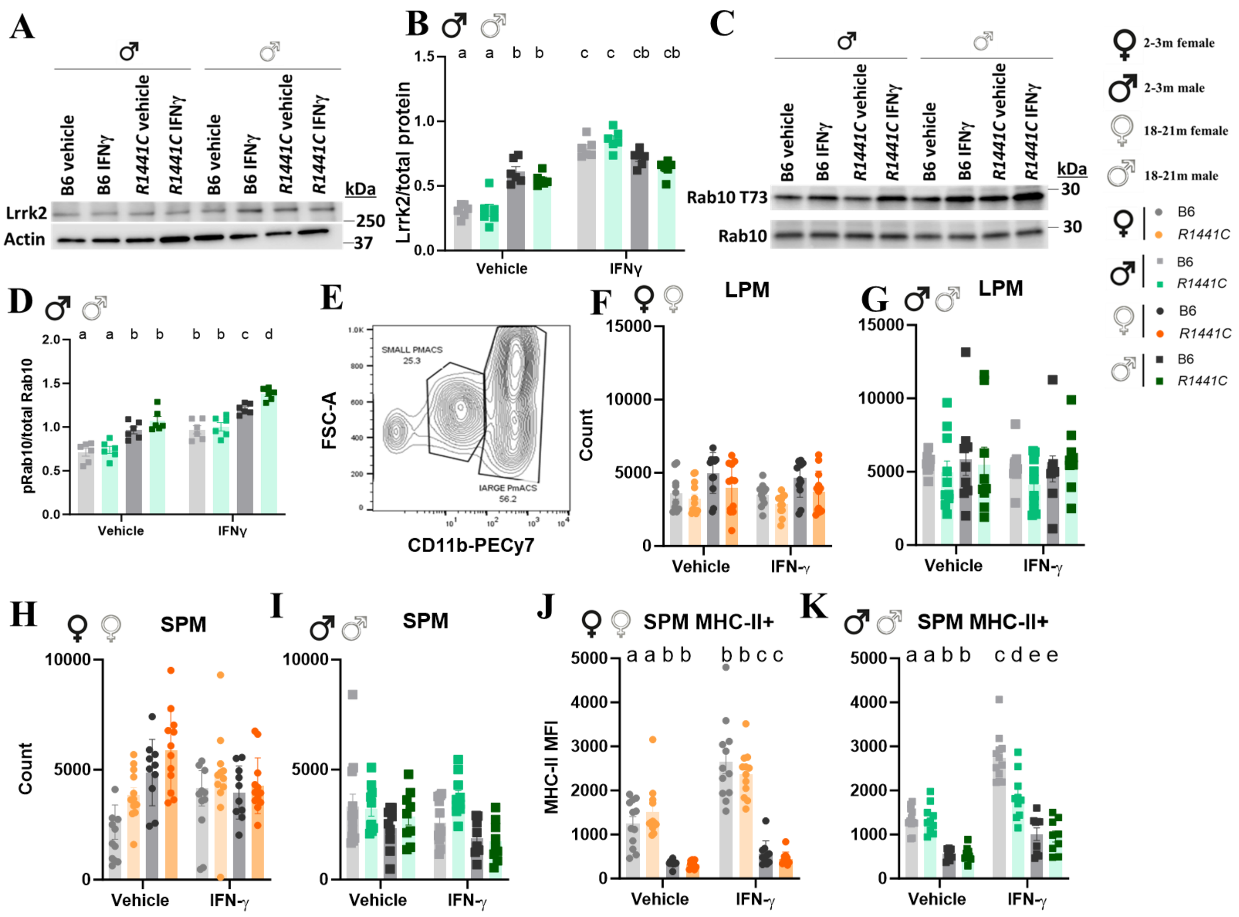


**Figure S1. Characterization of *ex vivo* pMac cultures.** pMacs from 2-3-month and 18-21-month-old female or male, B6 or *R1441C* mice were stimulated with 100U IFNγ for 18-hours. (A, B) Total Lrrk2 expression was assessed and normalized to β-Actin and quantified. Representative western blots shown. (C, D) T73 Rab10 protein expression was assessed and normalized to total Rab10 and quantified. Representative western blots shown. (E) Flow cytometry plot demonstrating differential expression of CD11b in LPMs and SPMs. (F-I). LPM and SPM counts were quantified via flow cytometry. (J, K) MHC-II expression on SPMs was quantified via flow cytometry. Bars represent mean +/- SEM (N = 8-12 mice). Three-way ANOVA, Bonferroni *post hoc*, groups sharing the same letters are not significantly different ($p > 0.05$) whilst groups displaying different letters are significantly different ($p < 0.05$).

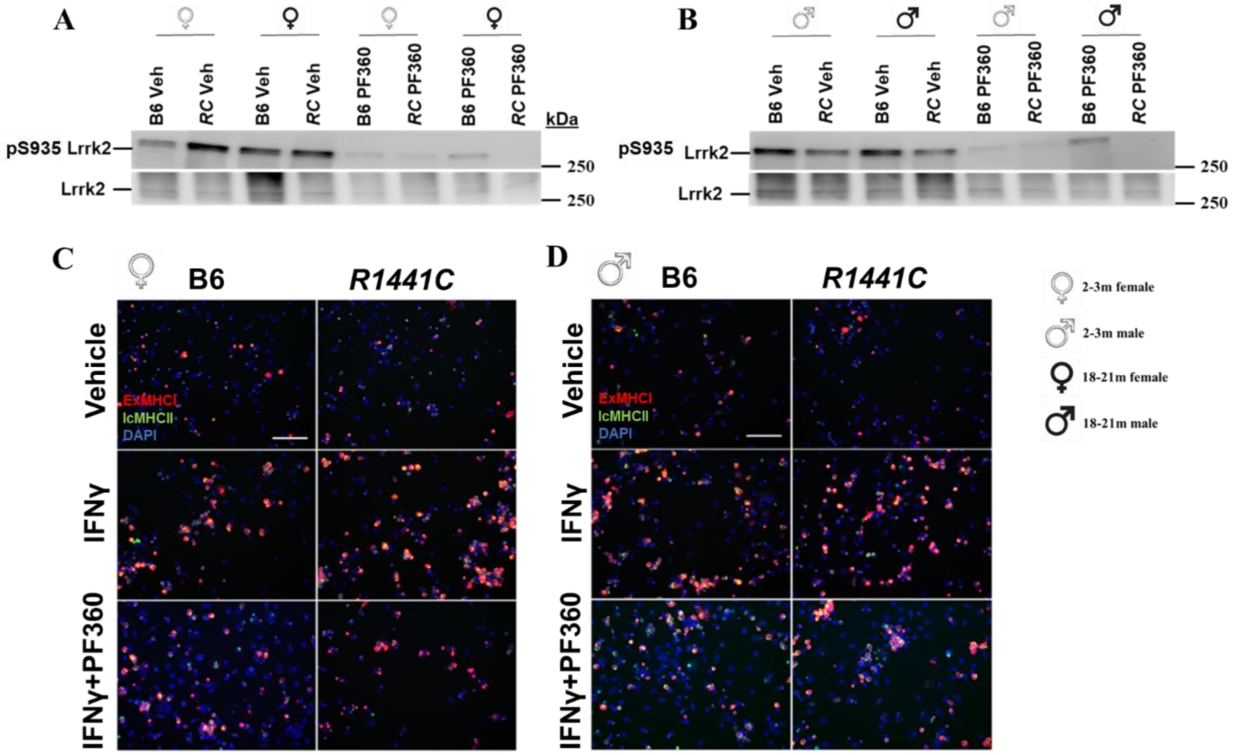


**Figure S2. *R1441C* mutation leads to age-dependent biphasic alteration in antigen presentation.** pMacs from 2-3-month and 18-21-month-old female or male, B6 or *R1441C* mice were treated with 100nM PF360 for 2-hours. (A, B) Representative western blots of S935 Lrrk2 and total Lrrk2 protein expression in pMacs. (C, D) pMacs were stained for intracellular and extracellular MHC-II. Scale bars, 100μM.

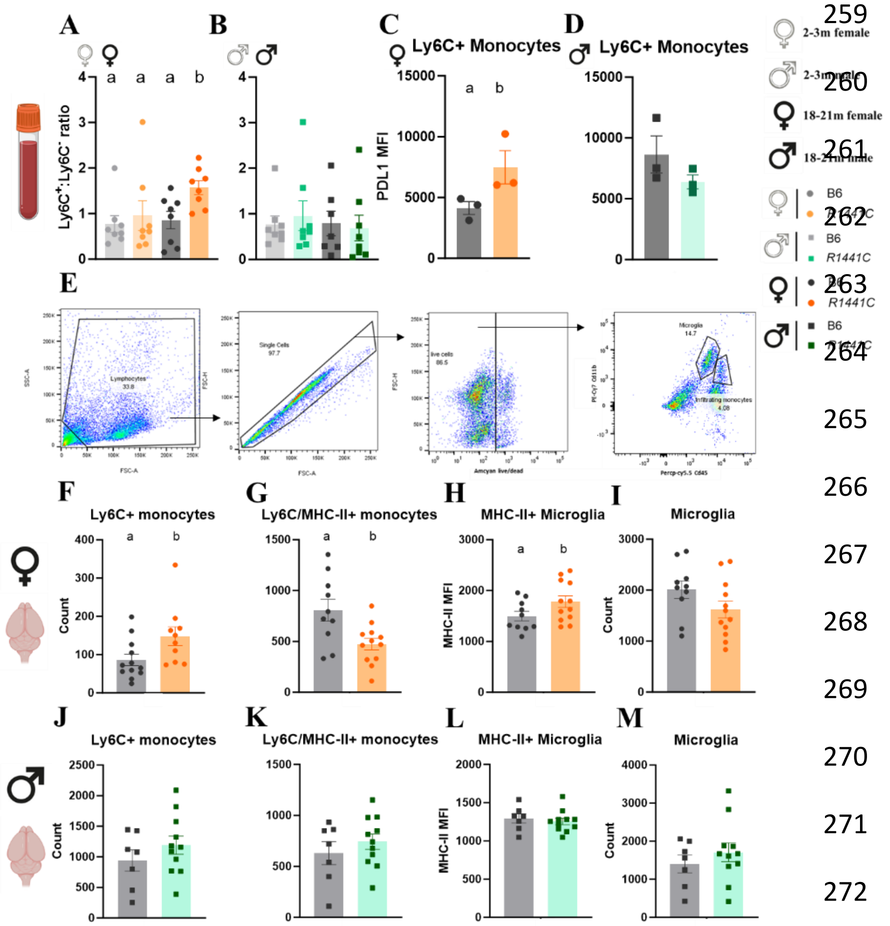


**Figure S3. The *R1441C* Lrrk2 mutation causes alterations of monocyte population and microglia activation in aged female mice**. (A, B) PBMCs from trunk blood of 2-3-month and 18-21-month-old female or male, B6 or *R1441C* mice were isolated and Ly6C+ and Ly6C- monocyte counts quantified via flow cytometry and ratios calculated. Bars represent mean +/- SEM (N = 3-10). One-way ANOVA, Bonferroni *post hoc*. (C, D) PBMCs from trunk blood of 18-21-month-old female or male, B6 or *R1441C* mice were isolated and PDL1 expression on Ly6C+ monocytes was quantified via flow cytometry. (E) Immune cells from the brain were isolated and microglia and infiltrating monocytes were identified via differential Cd11b and Cd45 expression. (F-M) Peripheral immune cells and microglia were isolated from the brains of 2-3-month and 18-21-month-old female or male, B6 or *R1441C* mice and cells counted and assessed for MHC-II expression. Bars represent mean +/- SEM (N = 3-10 mice). Student's T-test. Groups sharing the same letters are not significantly different ($p > 0.05$) whilst groups displaying different letters are significantly different ($p < 0.05$).

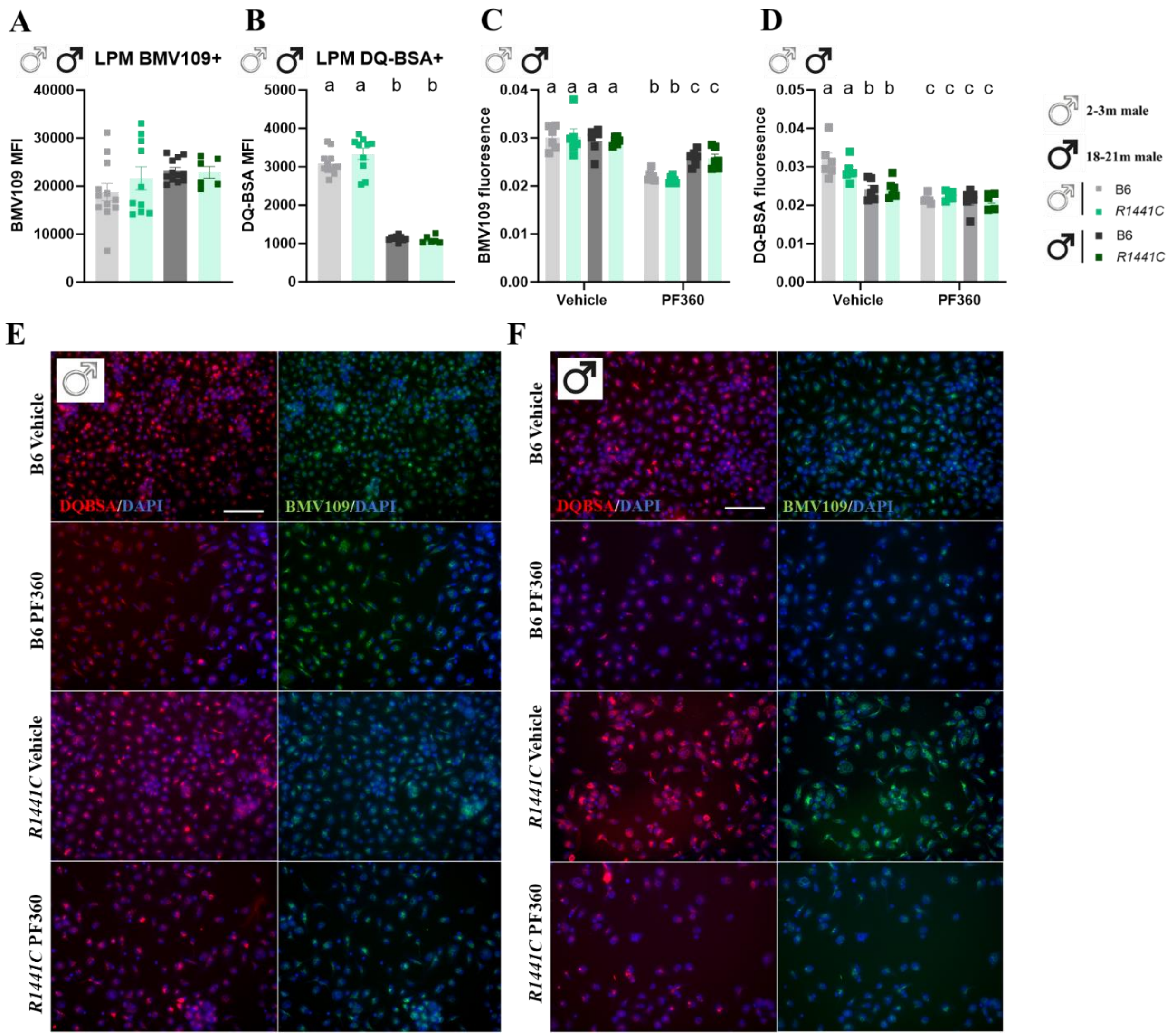


**Figure S4. *R1441C* mutation leads to age-dependent biphasic alteration in lysosomal activity**. pMacs from 2-3-month and 18-21-month-old male, B6 or *R1441C* mice were harvested and DQ-BSA and BMV109 MFI in LPMs quantified via flow cytometry (A, B). (C-F) pMacs were treated with vehicle or 100nM PF30 for 2 hours and DQ-BSA and BMV109 MFI was quantified via microscopy. Scale bars, 100µM. Bars represent mean +/- SEM (N = 6-12 mice). Two/Three-way ANOVA, Bonferroni *post hoc*. Groups sharing the same letters are not significantly different ($p > 0.05$) whilst groups displaying different letters are significantly different ($p < 0.05$).

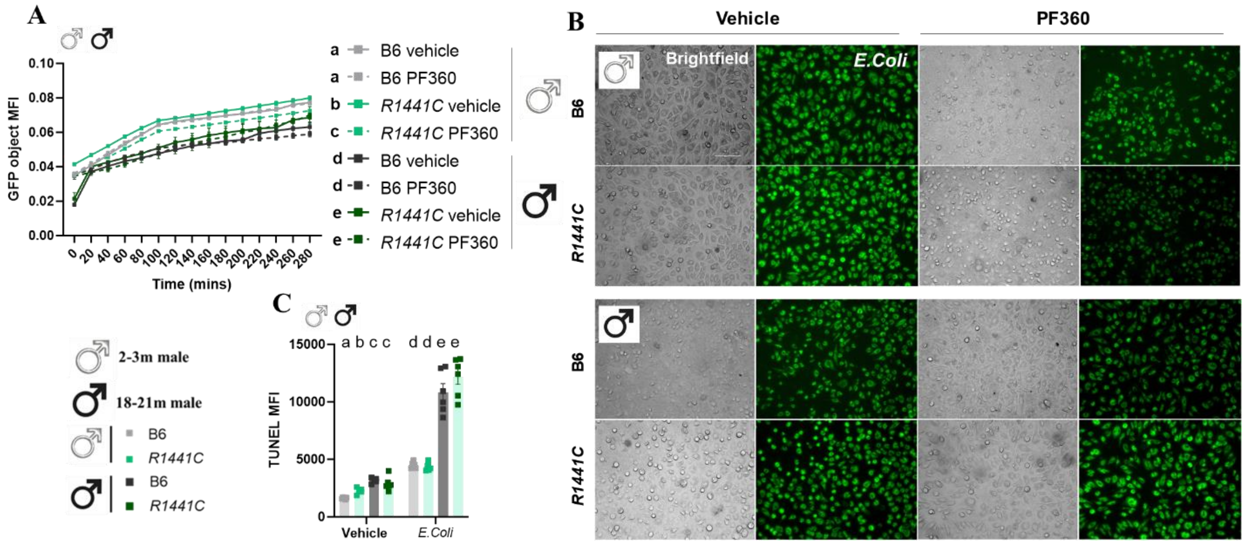


**Figure S5. *R1441C* Lrrk2 mutation alters pathogen uptake and pathogen-mediated DNA damage in age- and sex-dependent manner**. pMacs from 2-3-month and 18-21-month-old male, B6 or *R1441C* mice were incubated with 100µg of pHrodo *E. coli*. +/- 100nM PF360 (A, B) pMacs were imaged every 20 minutes for 5 hours and GFP MFI was measured. Scale bars, 100µM. Four-way ANOVA, Bonferroni *post hoc*. Main effects of groups are reported in each graph key. Groups sharing the same letters are not significantly different ($p > 0.05$) whilst groups displaying different letters are significantly different ($p < 0.05$). (C) TUNEL MFI was quantified in LPMs via flow cytometry. Bars represent mean +/- SEM (N = 6-12 mice). Three-way ANOVA, Bonferroni *post hoc*. Groups sharing the same letters are not significantly different ($p > 0.05$) whilst groups displaying different letters are significantly different ($p < 0.05$).

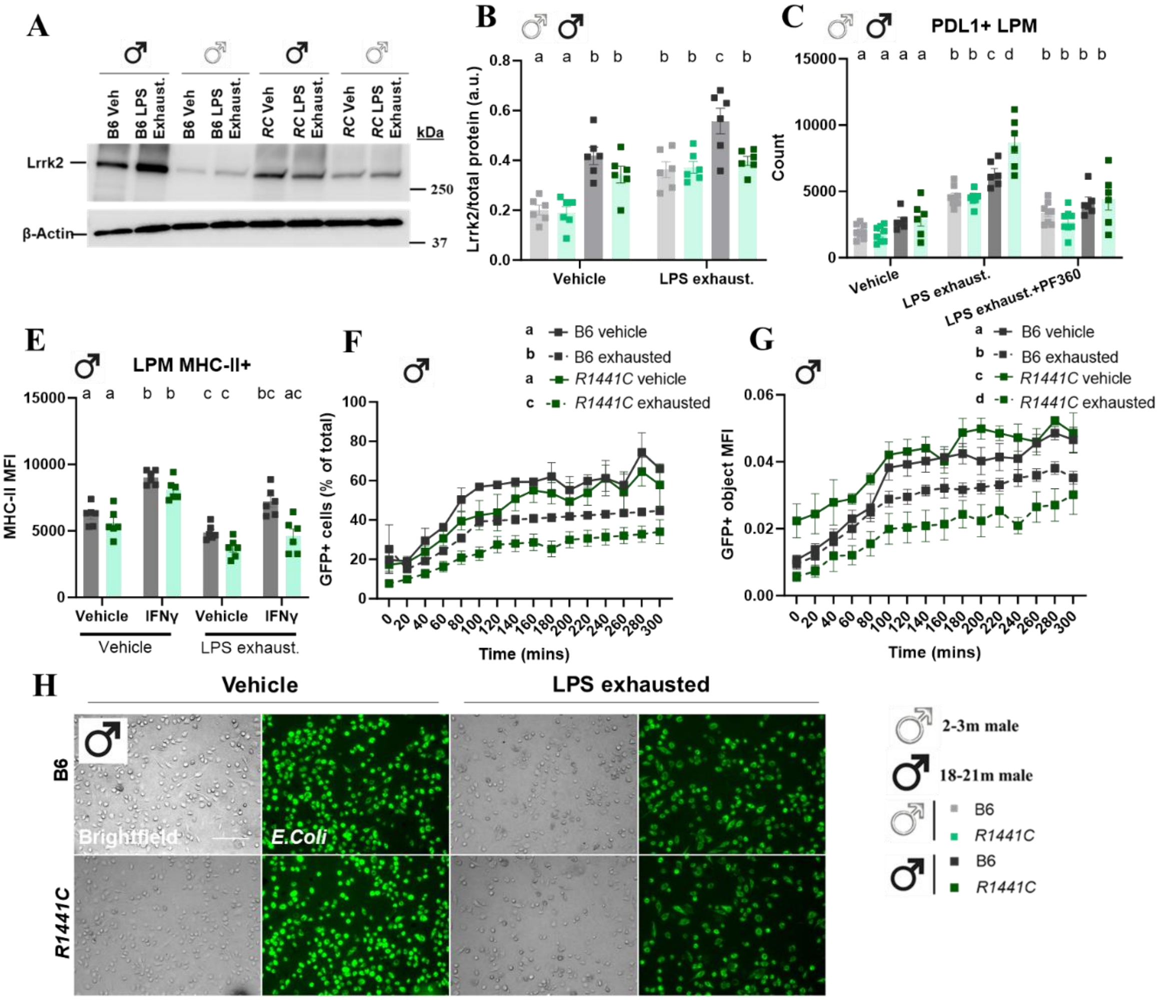


**Figure S6. *R1441C* Lrrk2 mutation induces immune cell exhaustion in a kinase-dependent manner.** pMacs from 2-3-month and 18-21-month-old male, B6 or *R1441C* mice were incubated with 100ng/mL of LPS for 5 days +/- 100nM PF360. (A, B) Lrrk2 protein expression were assessed and normalized to β-Actin and quantified. Representative western blots shown. (C) PDL1+ LPM count was quantified via flow cytometry. (E) After 5-day LPS or vehicle treatment, pMacs were stimulated with 100nG/mL of IFNγ for 18-hours and MHC-II MFI expression was quantified via flow cytometry. (F-H) After 5-day LPS or vehicle treatment, pMacs were incubated with 100μg of pHrodo *E. coli* pMacs were imaged every 20 minutes for 5 hours, and GFP+ cells (% of total) and GFP MFI was measured. Scale bars, 100μM. Bars represent mean +/- SEM (N = 6 mice). Three/Four-way ANOVA, Bonferroni *post hoc*. Main effects of groups are reported in each graph key. Groups sharing the same letters are not significantly different ($p > 0.05$) whilst groups displaying different letters are significantly different ($p < 0.05$).

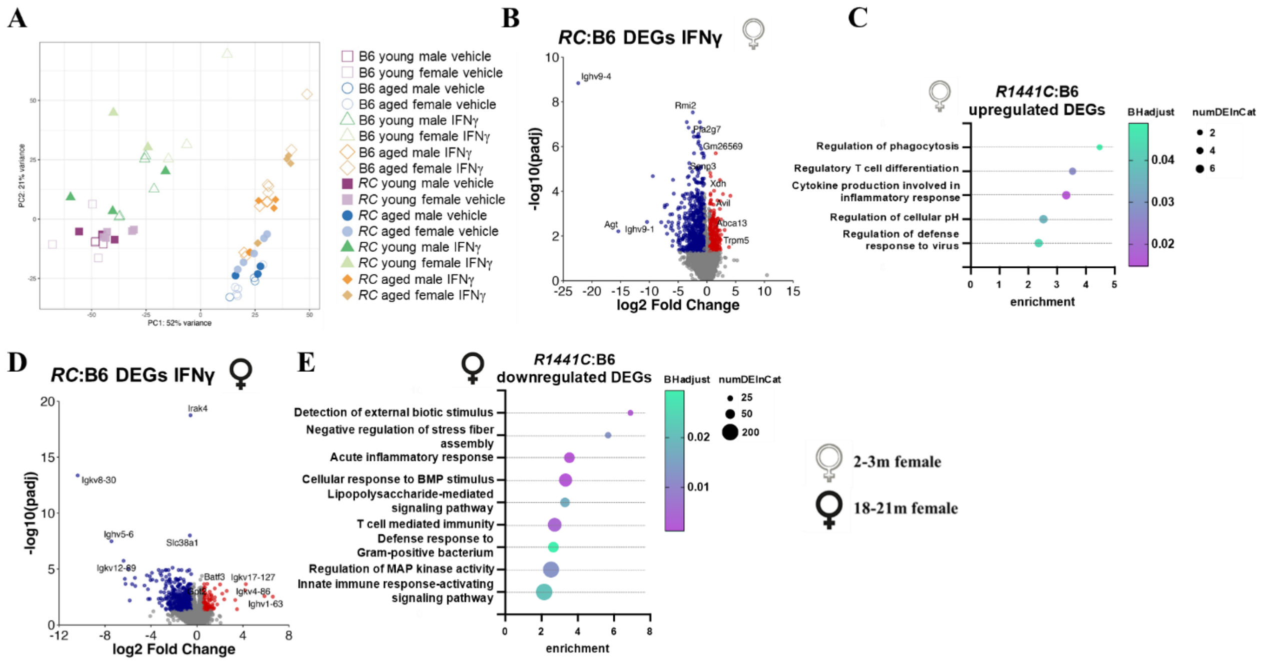


**Figure S7. Transcriptomic profiling reveals alterations in critical effector function pathways in *R1441C* macrophages in age- and sex- dependent manner.** (A). PCA plot generated from transcriptomic data from pMacs treated with vehicle or 100U IFNγ for 18h harvested from 2-3-month or18-21-month-old, female or male, B6 or *R1441C* mice. (B) Volcano plot of significant *R1441C*:B6 DEGs in IFNγ-treated pMacs from young females. (C) GO BP enrichment analysis identifies pathways in *R1441C*:B6 DEGs in IFNγ-treated pMacs from young female mice. (D) Volcano plot of significant *R1441C*:B6 DEGs in IFNγ-treated pMacs from aged females. (E) GO BP enrichment analysis identifies pathways enriched in which *R1441C*:B6 DEGs in IFNγ-treated pMacs from aged female mice. Volcano plots show genes with $\log_2$-transformed fold change > 0.5 and an adjusted p-value ≤ 0.05.

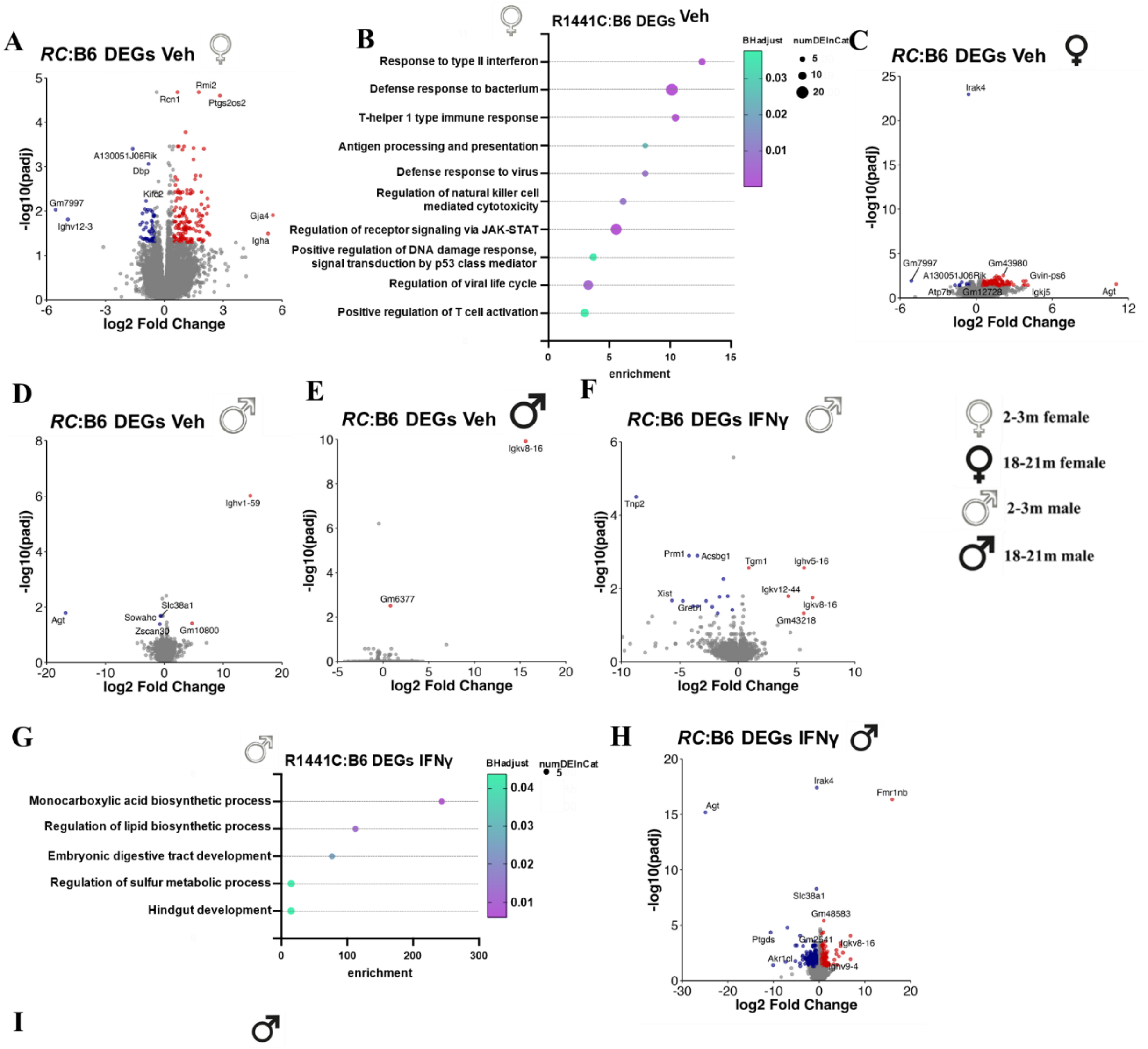


**Figure S8. *R1441C Lrrk2* mutation alters biological processes in macrophages in a sex- and age-dependent manner.** (A) Volcano plot of significant *R1441C*:B6 DEGs in vehicle-treated pMacs from young females. (B) GO BP enrichment analysis identifies pathways enriched in *R1441C*:B6 DEGs in vehicle-treated pMacs from young female mice. Volcano plots of significant *R1441C*:B6 DEGs in vehicle-treated pMacs from aged females (C), young males (D) and aged males (E). (F) Volcano plot of significant *R1441C*:B6 DEGs in IFNγ-treated pMacs from young males. (G) GO BP enrichment analysis identifies pathways enriched in which *R1441C*:B6 DEGs in IFNγ-treated pMacs from young male mice. (H) Volcano plot of significant *R1441C*:B6 DEGs in IFNγ-treated pMacs from aged males. (I) GO BP enrichment analysis identifies pathways enriched in *R1441C*:B6 DEGs in IFNγ-treated pMacs from aged male mice. Volcano plots show genes with $\log_2$-transformed fold change > 0.5 and an adjusted p-value ≤ 0.05.

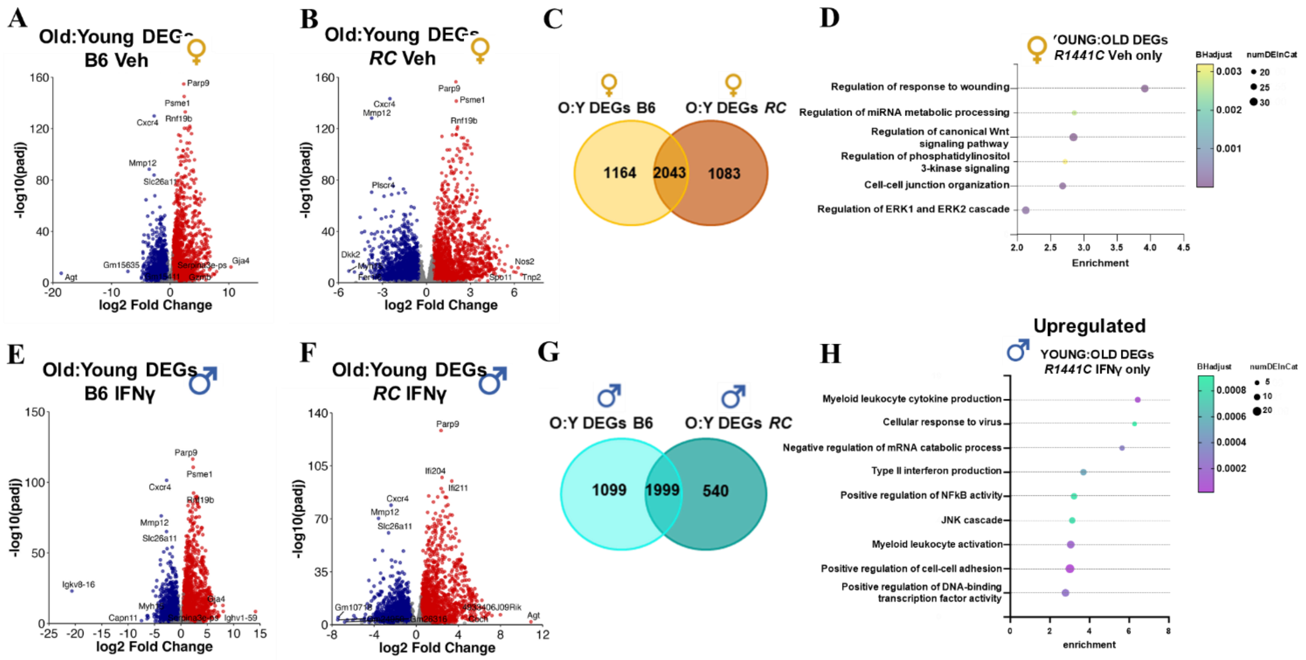


**Figure S9. *R1441C Lrrk2* mutation alters transcriptional changes associated with age in a sex-dependent manner.** (A, B, C). Significant Old:Young DEGs in vehicle-treated pMacs were counted and compared across B6 and *R1441C* female mice. Volcano plots show genes with fold change > 0.5 and an adjusted p-value ≤ 0.05. (D) GO BP enrichment analysis identifies pathways enriched in Old:Young DEGs seen only in *R1441C* female samples. (D, E, F). Significant Old:Young DEGs in IFNγ-treated pMacs were counted and compared across B6 and *R1441C* male mice. Volcano plots show genes with fold change > 0.5 and an adjusted p-value ≤ 0.05. (D) GO BP enrichment analysis identifies pathways enriched in Old:Young DEGs seen only in *R1441C* male samples.

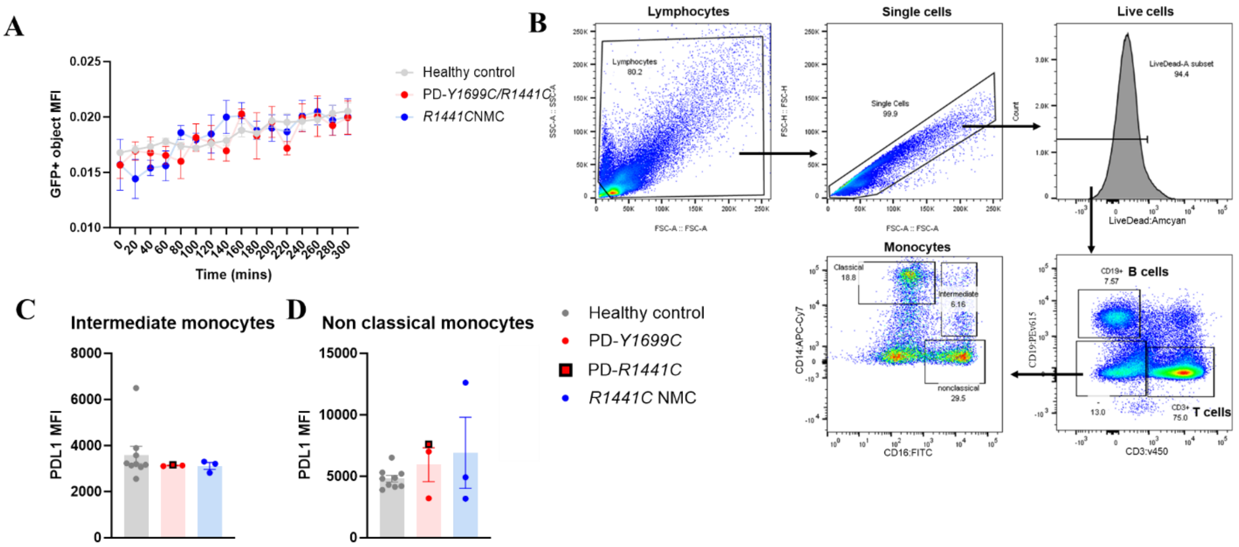


**Figure S10. *R1441C*- and *Y1699C*-LRRK2 monocyte derived macrophages exhibit hypophagocytosis, and increased markers for immune cell exhaustion.** (A) Macrophages were incubated with 100µg of pHrodo *E. coli* and imaged every 20 minutes for 5 hours, and GFP+ MFI was quantified. Three-way ANOVA, Bonferroni *post hoc*. (B) Flow gating strategy for human PBMCs. (C, D, E) PD1 MFI was quantified on T cells and monocyte subtypes via flow cytometry. One-way ANOVA, Bonferroni *post hoc*. Groups sharing the same letters are not significantly different ($p > 0.05$) whilst groups displaying different letters are significantly different ($p < 0.05$). (G) Monocyte-derived macrophages were stained for PD1 and microscopy images captured.

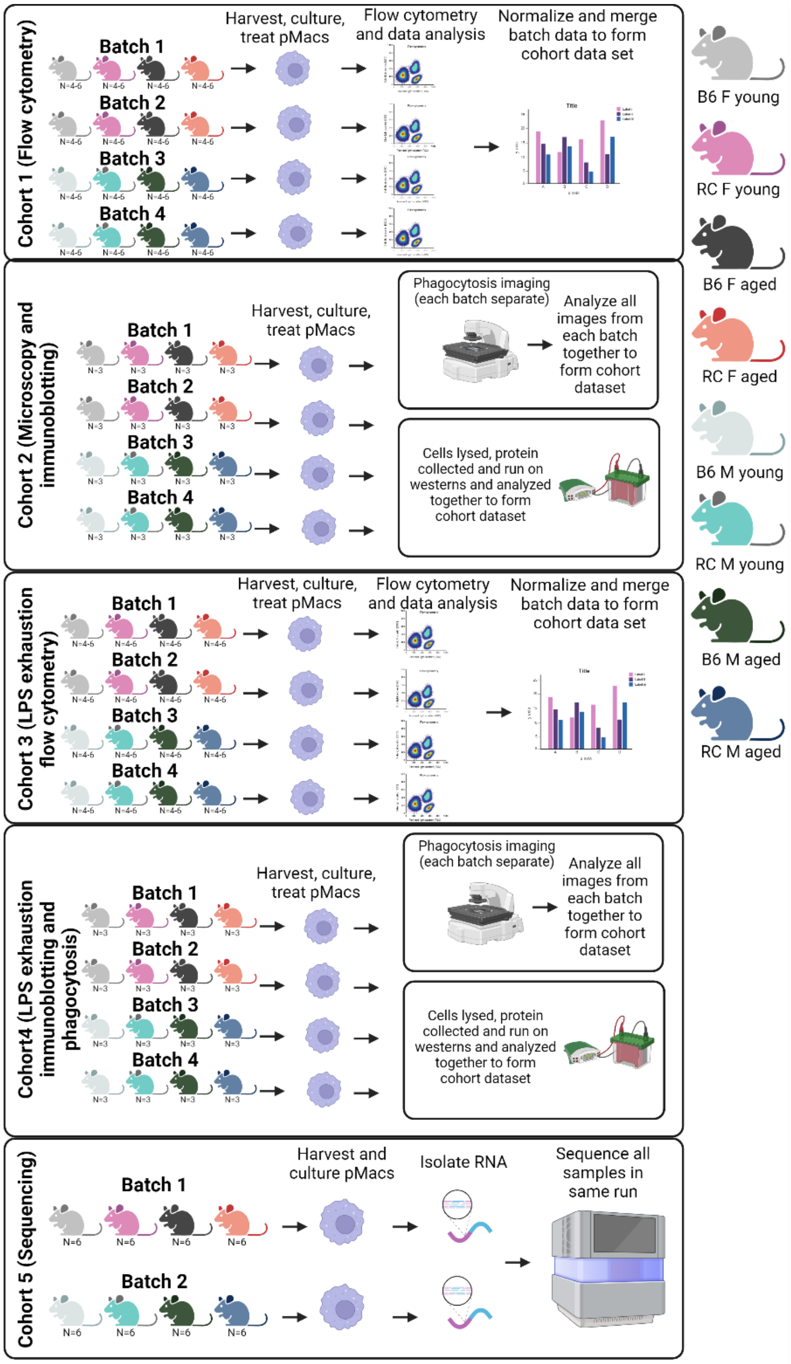


**Figure S11. Schematic of study design with mice**

**Supplementary tables**

**Table S1. Flow cytometry marker antibody panel (mouse)**

| Target | Conjugate | Antibody Cat# | Dilution | Company | RRID |
|---|---|---|---|---|---|
| **CD11b** | PE-Cy7 | 101216 | 1:100 | Biolegend | AB_312798 |
| **MHC-II** | APC-Cy7 | 107628 | 1:100 | Biolegend | AB_10612741 |
| **Live/dead stain** | Amcyan | 130113144 | 1:2000 | Fisher | AB_2725972 |
| **FcR** | - | 422301 | 1:100 | Biolegend | AB_2818986 |
| **eBioYAe** | FITC | 11-5741-82 | 1:100 | ThermoFisher | AB_996692 |

**Table S2. Antibodies for immunoblotting**

| Target | Antibody Cat# | Dilution | Company | RRID |
|---|---|---|---|---|
| **LRRK2** | ab133474 | 1:1000 | Abcam | AB_2713963 |
| **LRRK2 pS935** | ab133450 | 1:1000 | Abcam | AB_2732035 |
| **Rab10** | MABN730 | 1:1000 | Millipore | NA |
| **Rab10 pT73** | ab241060 | 1:1000 | Abcam | NA |
| **B-actin** | AM4302 | 1:5000 | Thermo | AB_2536382 |
| **Revert™ 700 Total Protein Stain** | 926-11011 | NA | Licor | NA |
| **Goat Anti-Rabbit IgG HRP** | 111035144 | 1:5000 | Jackson lab | AB_2307391 |
| **Goat Anti-Mouse IgG HRP** | 115-036-072 | 1:5000 | Jackson lab | AB_2338525 |

**Table S3. Patient demographics**

| Group | N | Mean Age (years) | Mean Age at diagnosis (years) | Sex (Male/Female) |
|---|---|---|---|---|
| **Healthy controls** | 9 | 40.8 | - | (5/4) |
| ***R1441C*-PD** | 1 | 42 | 27 | (1/0) |
| ***Y1699C*-PD** | 2 | 63.5 | 56.5 | (2/0) |

| | | | | |
|---|---|---|---|---|
| ***R1441C*-NMC** | 3 | 30.5 | - | (3/0) |

**Table S4. Flow cytometry marker antibody panel (human)**

| Target | Conjugate | Antibody Cat# | Dilution | Company | RRID |
|---|---|---|---|---|---|
| **CD3** | V450 | 560365 | 1:20 | Biolegend | AB_1645570 |
| **CD16** | FITC | 302006 | 1:20 | Biolegend | AB_314206 |
| **CD14** | APC-v770 | 130-098-076 | 1:20 | Miltenyi | AB_2660181 |
| **CD19** | PE-615 | 130-108-388 | 1:20 | Miltenyi | AB_2661292 |
| **PD1** | APC | 563741 | 1:20 | BD Bioscience | AB_2738399 |
| **FcR** | - | 422301 | 1:20 | Biolegend | AB_2818986 |
| **Live/dead stain** | Amcyan | 130113144 | 1:2000 | Fisher | AB_2725972 |

### References only cited in SM
